# Bioactive Hydrogel Microcapsules for Guiding Stem Cell Fate Decisions by Release and Reloading of Growth Factors

**DOI:** 10.1101/2021.09.21.461208

**Authors:** Kihak Gwon, Hye Jin Hong, Alan M. Gonzalez-Suarez, Michael Q. Slama, Daheui Choi, Jinkee Hong, Harihara Baskaran, Gulnaz Stybayeva, Quinn P. Peterson, Alexander Revzin

## Abstract

Human pluripotent stem cells (hPSC) hold considerable promise as a source of adult cells for treatment of diseases ranging from diabetes to liver failure. Some of the challenges that limit the clinical/translational impact of hPSCs are high cost and difficulty in scaling-up of existing differentiation protocols. In this paper, we sought to address these challenges through the development of bioactive microcapsules. A co-axial flow focusing microfluidic device was used to encapsulate hPSCs in microcapsules comprised of an aqueous core and a hydrogel shell. Importantly, the shell contained heparin moieties for growth factor (GF) binding and release. The aqueous core enabled rapid aggregation of hPSCs into 3D spheroids while the bioactive hydrogel shell was used to load inductive cues driving pluripotency maintenance and endodermal differentiation. Specifically, we demonstrated that one-time 1h long loading of pluripotency signals, fibroblast growth factor (FGF)-2 and transforming growth factor (TGF)-β1, into bioactive microcapsules was sufficient to induce and maintain pluripotency of hPSCs over the course of 5 days at levels similar to or better than a standard protocol with soluble GFs. Furthermore, stem cell-carrying microcapsules that previously contained pluripotency signals could be reloaded with an endodermal cue, Nodal, resulting in higher levels of endodermal markers compared to stem cells differentiated in a standard protocol. Overall, bioactive heparin-containing core-shell microcapsules decreased GF usage five-fold while improving stem cell phenotype and are well suited for 3D cultivation of hPSCs.

## 1. Introduction

Human pluripotent stem cells (hPSC) proliferate indefinitely and may be differentiated into any adult cell type [1–3]. This makes hPSCs an excellent cell source for regenerative medicine and tissue engineering applications [4, 5]. hPSC maintenance and differentiation protocols rely on inductive cues, most often growth factors (GFs), that are added into culture media daily in ng/mL quantities in a prescribed schedule over the course of days and weeks [6, 7]. While well-established and characterized, such differentiation protocols require significant amounts of recombinant GFs, which make it costly to scale up to larger volume stem cell cultures. The objective of this paper was to explore maintenance and differentiation of hPSCs in bioactive microcapsules where inductive GFs may be loaded once and then released locally to the encapsulated hPSCs over the course of several days.

There are a number of recent reports describing the use of biomaterial scaffolds for expansion and differentiation of hPSCs in a 3D format [8–11]. However, these strategies employed macro-scale scaffolds or extruded biomaterial filaments carrying stem cells. We and others have focused on developing microcapsules for 3D cultivation of hPSCs [12, 13]. Encapsulated hPSC spheroids are easy to handle and may be dispensed into microtiter plates for culture optimization, disease modeling or therapy testing experiments. Microcapsules may also be used as carriers for scalable cultivation of stem cells in suspension cultures. Early on, our lab focused on encapsulating stem cells in solid gel microparticles but found that while mouse embryonic stem cells (mESCs) survived and thrived in such microcapsules, hESCs did not fare as well [14]. We reasoned that hESCs may be more dependent on cell-cell contacts than mESCs, and may benefit from microcapsules that contained aqueous environment where cells could rapidly aggregate. We fabricated core-shell microcapsules comprised of poly (ethylene glycol) (PEG) hydrogel shell and an aqueous core [15], and demonstrated that hPSCs may be successfully maintained and differentiated in such core-shell microcapsules [12]. This encapsulation strategy was shown to have several benefits: 1) individual hPSCs confined in an aqueous core rapidly aggregated and formed spheroids, 2) the size of spheroids could be controlled precisely by the initial cell density, and 3) the hydrogel shell of the microcapsule offered protection against shear stress during cultures in a stirred bioreactor. However, this past study employed biologically inert microcapsules composed of PEG and relied on soluble inductive cues for cultivation and differentiation of hPSCs. In the present study, we explored the use of heparin-containing core-shell microcapsules for loading and release of GFs that drive fate selection of encapsulated hPSC spheroids.

There is strong and long-standing interest in the tissue engineering community to develop scaffolds comprised of extracellular matrix (ECM) elements capable of sequestration and release of GFs. A number of scaffolds incorporating peptides, proteins or polysaccharides have been described in the literature [16, 17]. Heparin is a naturally occurring polysaccharide and an ECM component that has been used to design biomaterials capable of sequestration and release of GFs [18, 19]. The sequestration occurs *via* a secondary bond formation between heparin and heparin-binding domains expressed on multiple GFs. First described by Sakiyama-Elbert and Hubbell [20], the concept of heparin-containing hydrogels was refined by Tae and Stayton who covalently incorporated functionalized heparin into a PEG hydrogel network [21, 22]. Over the years, heparin-containing hydrogels have been used for loading and controlled release of a number of GFs including acidic and basic fibroblast growth factors (aFGF and bFGF), hepatocyte growth factor (HGF), vascular endothelial growth factor (VEGF), transforming growth factor (TGF)-β1, and Nodal [14, 18, 21–23].

Beyond loading and release of GFs, a number of studies have shown that heparin-containing scaffolds contribute to improved proliferation, differentiation, and function of entrapped cells [24–26]. We have previously employed droplet microfluidics to fabricate bioactive heparin-containing hydrogel microparticles and demonstrated successful encapsulation and endodermal differentiation of mESCs. Loading of endodermal signaling molecule, Nodal, into microparticles and differentiation of mESCs were also demonstrated. However, solid hydrogel microparticles used by us in this past study proved suboptimal for encapsulating human (h) ESCs, which are less proliferative and more reliant on reestablishing cell-cell contacts than mouse counterparts [14].

The objective of this study was to develop hydrogel microcapsules incorporating heparin moieties for GF loading and an aqueous core for hESC aggregation and spheroid formation. To the best of our knowledge, core-shell microcapsules with such properties have not been reported to date. We characterized release profiles of pluripotency signals, FGF-2 and TGF-β1, and endodermal cue, Nodal, from bioactive microcapsules. We also demonstrated that pluripotency and endodermal signals may be loaded and released in a sequential manner from the same microcapsules (see Scheme 1). Compared to standard protocols employing soluble GFs, one-time loading and sustained release of inductive cues allowed us to decrease the amount of GF required for pluripotency maintenance and endodermal differentiation by a factor of 5. Beyond cost reduction, local release of GFs from bioactive microcapsules resulted in hPSCs expressing higher levels of pluripotency and endodermal markers compared to standard protocols.

**Scheme 1.**
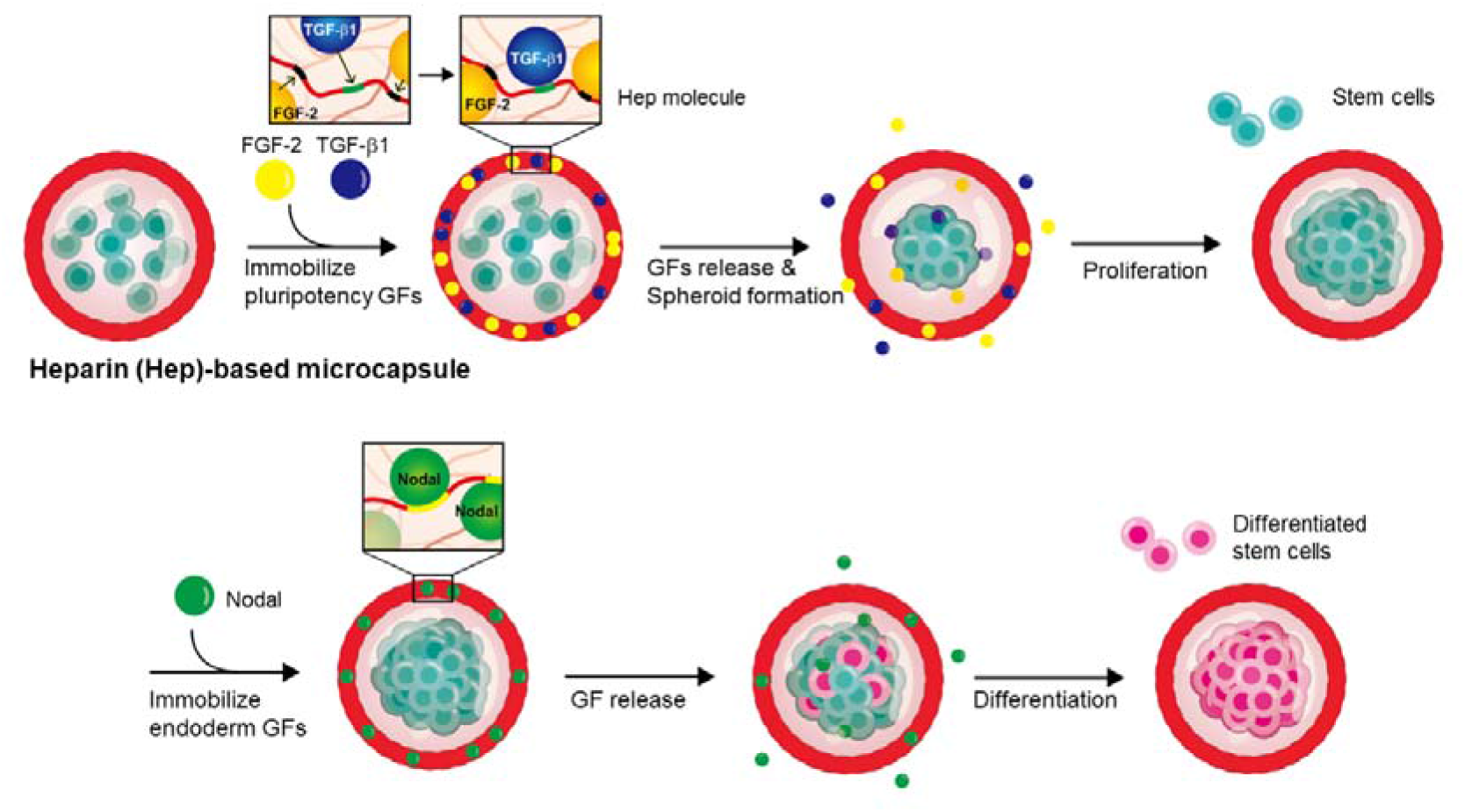
Bioactive microcapsules for sequential loading and release of stem cell inductive cues. TGF-β1 and FGF-2, GFs required for maintenance of pluripotency, are loaded into hPSC-carrying microcapsules first. Five days later, after pluripotency cues have been delivered to hPSCs, microcapsules are reloaded with endodermal signal, Nodal, to start the differentiation process.

## 2. Materials and Methods

### 2.1. Synthesis of methacrylated heparin (Hep-MA)

Heparin was functionalized with methacrylate groups as described previously [14]. Briefly, 200 mg of sodium heparin (12 kDa; Smithfield Bioscience, Cincinnati, OH, USA) was first dissolved in 10 mL filtered distilled water (DW) to make a 2% (w/v) heparin solution. Then the solution was reacted with methacrylic anhydride (MA) in 10-fold molar excess (Sigma-Aldrich, St. Louis, MO, USA) for 12 h in the dark at 4 □ while maintaining the pH between 8 to 11 using 5 N or 1 N NaOH. The final product (Hep-MA) was precipitated in cold absolute ethanol with 10-fold volume (Sigma-Aldrich). After centrifugation at 5000 g for 5 min at 4 □, the supernatant was removed, and the precipitant was re-dissolved in 10 mL DW. To remove any unreacted reagents, Hep-MA was purified by dialysis against DW using a 3.5 kDa MW cut-off dialysis membrane (Spectrum Laboratories, Rancho Dominguez, CA, USA) for 3 days. The resultant solution was freeze-dried (Sentry 2.0, SP Scientific, Warminster, PA, USA) for 3 days and stored at −20 □ for later use. The degree of methacrylation was determined by protein nuclear magnetic resonance (^1^H NMR) spectroscopy (DMX 360; Bruker, Billerica, MA, USA) by dissolving 20 mg of Hep-MA in 600 μL deuterium oxide (D_2_O) (Sigma-Aldrich).

### 2.2. Fabrication of microfluidic encapsulation devices

Co-axial flow-focusing devices were fabricated based on a previously described protocol [12]. First, we fabricated two master molds on 4-inch silicone (Si) wafers using a multi-step photoresist patterning process described in Fig. S1A. One Si wafer contained the original design while the other had the mirror image of the original. The wafers were processed in three spin-coating and photolithography steps to generate SU-8 structures with the following heights: 1) 60 μm for core stream, 2) 100 μm for shell stream, and 3) 150 μm for oil streams. The master wafers were later replicated into polydimethylsiloxane (PDMS) slabs with imprinted channel architecture. Subsequently, the two PDMS slabs were aligned under a stereoscope and bonded to fabricate an encapsulation device. Each completed device had channels of the following heights: 1) core – 120 μm, 2) shell – 200 μm and 3) oil – 300 μm. Master mold fabrication and PDMS replication processes are described in detail below.

To fabricate core channels of the encapsulation device, SU-8 2025 (Kayaku Advanced Materials, Westborough, MA, USA) was spin-coated on 4-inch silicon wafers (University Wafer, USA) to a thickness of 60 μm (1300 rpm) and soft baked (3 min at 65 □, 15 min at 95 □). The SU-8 layer was then exposed using a direct-write photolithography tool (μPG 101; Heidelberg Instrument, Woburn, MA, USA) that contains a 70 mW UV laser (375 nm) where energy can be modulated. The first layer was exposed using an energy of 50 mW at 100%. The exposed SU-8 underwent a post exposure bake for 2 min at 65 □, followed by 10 min at 95 □. The wafer was submerged for ~3 min in SU-8 developer (Kayaku Advanced Materials) to remove unreacted photoresist and placed on a hot plate for 20 min at 160 □ to (hard bake) improve resist adhesion to substrate. After letting the wafer cool down to 25 □, it was spin-coated with SU-8 2050 photoresist to a thickness of 100 μm (1400 rpm) in order to generate the second layer containing shell structures of the device. After spin-coating, the wafer was soft baked for 5 min at 65 □ and 30 min at 95 □ then exposed using the μPG 101 (50 mW, 100%, 2 consecutive exposures). Exposure was followed by a post exposure bake for 5 min at 65 □ and 10 min at 95 □. Development of SU-8 and hard bake was carried out in a similar manner to that described for the first layer. Next, we spin-coated SU-8 2050 to a thickness of 150 μm (1000 rpm) for the oil channel network of the device. Soft bake and exposure parameters were identical to those previously described for layer two. Post exposure bake was carried out for 5 min at 65 □ and 35 min at 95 □, followed by developing and hard bake steps. After depositing the third SU-8 layer, the molds were treated in vapors of chlorotrimethylsilane (Sigma-Aldrich) for 1 h inside a covered 150 mm glass dish to minimize adhesion of PDMS to the master molds.

After both master molds were prepared, top and bottom parts of the device were generated by the following procedure. First, a master mold was placed on a plastic Petri dish (150 mm diameter) and a PDMS (Sylgard 184 silicone elastomer kit; Dow Corning, Midland, MI, USA) base was mixed with a curing agent in a 10:1 ratio, then the mixture was poured onto both master molds. PDMS prepolymer was degassed in a desiccator under vacuum for 30 min and cured in a convection oven at 80 □ for 2 h. Cured PDMS pieces were cut along the outline of the device, then punched by 14 G and 15 G needles for inlets and outlet, respectively. The PDMS pieces were treated with oxygen plasma (G-500 plasma cleaning system; Yield Engineering Systems, Livermore, CA, USA) for 20 s. Then the two PDMS pieces were aligned manually under a stereoscope and incubated in an oven at 80 □ for 2 h for complete bonding.

In addition to the encapsulation device described above, we also fabricated a filter device that is connected upstream of the encapsulation device. This device captures cell aggregates, improving cell suspension homogeneity and preventing clogging in core microchannel. A master mold for the filter device was fabricated on a 4-inch Si wafer using a single spincoating and photolithography patterning step as described in Fig. S1B. A master mold was then replicated using soft-lithography and PDMS as described in the preceding paragraph. PDMS pieces with imbedded channel architecture were bonded to glass substrates using oxygen plasma. A filter device (see Fig. S2 for device design) consisted of a single 50 μm tall chamber that was wider at the inlet area and narrower at the outlet. An array of triangular-shaped posts covered the whole chamber with spacing between posts decreasing as chamber width decreased toward the outlet. Triangle posts were 200 μm per side with pitch ranging from 400 (at the inlet) to 30 μm (at the outlet). The spacing of the posts was designed to ensure that cell aggregates entering the filter device were trapped, but also allowing breaking of loose aggregates as they pass through the device. Velocity and shear stress profiles in this device were modeled using COMSOL (Fig. S2).

### 2.3. Operation of microfluidic devices and fabrication of core-shell microcapsules

The coaxial flow-focusing microfluidic devices were loaded with 4 different solutions for core-shell microcapsule generation [12, 15]: 1) the core solution consisted of 8% (w/v) PEG (35 kDa; Sigma-Aldrich) and 17% (v/v) Optiprep densifier (STEMCELL Technologies, Vancouver, Canada) dissolved in DMEM/F12 medium (Gibco, Thermo Fisher Scientific, Waltham, MA, USA); 2) shell solution with 4% (w/v) Hep-MA, 8% (w/v) 4-arm PEG-maleimide (PEG4MAL) (10 kDa; Laysan Inc., Arab, AL, USA), and 15 mM triethanolamine (TEA; Sigma-Aldrich) dissolved in phosphate buffered saline (PBS); 3) the shielding oil consisted of 0.5% (v/v) Span-80 (Sigma-Aldrich) surfactant dispersed in mineral oil (Sigma-Aldrich); 4) crosslinking emulsifier containing 60 mM dithiothreitol (DTT; Sigma-Aldrich) dispersed in mineral oil with 3% Span-80. The crosslinking emulsion was sonicated in an ultrasonic bath at 20 □ for 1 h before use.

All four solutions were loaded into disposable syringes (Thermo Fisher Scientific), which were then assembled with 27 G needles. The solutions were all injected into the devices through Micro Medical Tubing (0.015” ID x 0.043” OD; Scientific Commodities, Inc., Lake Havasu City, AZ, USA) by syringe pumps (Harvard Apparatus, Holliston, MA, USA) at the following rates: core (4 μL/min), shell (4 μL/min), shielding oil (50 μL/min), and crosslinking emulsion (60 μL/min). In this study, the concentration of the PEG4MAL and Hep-MA was fixed to 8% (w/v) and 4% (w/v), respectively.

### 2.4. Characterization of bioactive core-shell microcapsules

#### 2.4.1. Encapsulation of fluorescent microbeads

The fluorescent yellow microbeads (10.2 μm; Spherotech, Lake Forest, IL, USA) were incubated with Pierce protein-free blocking buffer (Thermo Fisher Scientific) for 2 h at 4 □ and washed with PBS. For encapsulation, the microbeads were dispersed in the core solution to a final concentration of 1% (w/v) and rhodamine-PEG-SH or FITC-PEG-SH (5 kDa; Nanocs, Boston, MA, USA) was mixed in the shell solution in a 1:1000 molar ratio [12]. The bright-field and fluorescent images of microcapsules were obtained with an inverted fluorescence microscope (IX83; Olympus, Center Valley, PA, USA).

#### 2.4.2. SEM imaging of microcapsules

The core-shell microcapsules were imaged using Field-Emission Scanning Electron Microscopy (FE-SEM; Hitachi S-4700, Hitachi High Technologies America, Inc., Schaumburg, IL, USA) at 30 kV accelerating voltage. The core-shell microcapsules were fully swollen with DW for 24 h and frozen at −80 □ before going through lyophilization for 24 h [27]. Then, the lyophilized microcapsules were coated with gold-palladium (Au/Pd) for 30 s prior to SEM analysis.

#### 2.4.3. Diffusivity of microcapsules

Diffusivity properties of the microcapsule shells were characterized using fluorescent dextrans of different molecular weights. Microcapsules (~2000) were submerged in a well of a 6-well plate containing 10 μM TRITC-labeled dextran of different molecular weights: 4, 20, 70, and 2000 kDa (Sigma-Aldrich). After loading TRITC-dextran in the capsules by soaking (typically 3 h), microcapsules were transferred into a different well of a 6-well plate filled with 2 mL of fresh 1x PBS. A 6-well plate was placed on a fluorescence microscope and was imaged every 30 min to quantify changes in the fluorescence intensity inside microcapsules. The average fluorescent intensity was determined using an Image J software. The intensity was normalized to the initial intensity of the microcapsules and the time-dependent normalized intensity values were analyzed using an unsteady radial transport equation for the core. The transport in the shell was modeled using a permeability parameter. The equations were non-dimensionalized (see *Supplemental Information*) and solved using pde function in MATLAB^®^. The dimensionless permeability parameter was estimated using fmincon function in MATLAB^®^. Sum of squared differences between estimated and experimental values was minimized by adjusting the permeability parameter. Diffusivity values of dextrans in the shell were evaluated from the permeability parameter from the shell dimensions.

#### 2.4.4. Incorporation of heparin into microcapsules

The core-shell microcapsules were stained with toluidine blue O to verify heparin incorporation [14, 28]. The microcapsules were incubated in 10 mg/mL toluidine blue O (Sigma-Aldrich) solution for 30 min, with negatively charged heparin molecules in the hydrogel interacting with positively charged toluidine blue O *via* electrostatic interaction. Stained microcapsules appeared purple when imaged in bright-field mode (Zeiss Stemi DV4 Stereo Microscope; Carl Zeiss, Jena, Germany). The microcapsules fabricated in three different conditions were stained with toluidine blue O: 1) 4% (w/v) Hep-MA and 8% (w/v) PEG4MAL, 2) 4% (w/v) Hep and 8% (w/v) PEG4MAL, and 3) 4% (w/v) dimethacrylated PEG (PEG-DMA) and 8% (w/v) PEG4MAL.

### 2.5. Growth factor loading and release studies

For GF loading, 3600 microcapsules with or without heparin were placed into a well of a 6-well plate containing 2 mL of the following GF in 1x PBS: 1) FGF-2 (100 ng/mL), 2) TGF-β1 (2 ng/mL), and 3) Nodal (200 ng/mL). After 1 h at 37 □, the capsules were collected using a 100 μm cell strainer (Cardinal Health, Dublin, OH, USA) and the residual solution was analyzed by enzyme-linked immunosorbent assay (ELISA). To establish a GF release profile, 3600 microcapsules were placed into a well of a 6-well plate containing 2 mL of 1x PBS. At different time points (4 hours, 1, 3, 5 and 7 days), the entire solution containing released GF was exchanged with fresh PBS. The collected samples were analyzed for GF by ELISA. ELISA kits for FGF-2 and TGF-β1 were purchased from R&D Systems (Minneapolis, MN, USA) while the ELISA kit for Nodal was obtained from Abcam (Cambridge, MA, USA). ELISAs were performed as per manufacturers’ instructions.

### 2.6. Microencapsulation and cultivation of hPSCs

HUES-8 cells (hESC line) were maintained as spheroids in 30 mL spinner flasks (ABLE Biott, Beltsville, MD) as described previously [12]. The cells were suspended in mTeSR medium (STEMCELL Technologies) with 10 μM ROCK Inhibitor (Y27632; STEMCELL Technologies) in a concentration of 5 x 10^5^ cells/mL. The spinner flasks were placed on a stirring plate at the speed of 70 rpm inside the humidified incubator at 37 □ and 5% CO_2_ (Thermo Fisher Scientific). After 48 h, the medium was replaced with fresh mTeSR medium without Y27632. The cells were passaged every 72 h by dissociating to single cells using Accutase (Sigma-Aldrich) and re-suspended in fresh mTeSR medium with Y27632.

HUES-8 cells were collected by centrifugation (300 g for 5 min) and re-suspended at a concentration of 5 x 10^7^ cells/mL with the viscous core solution, which consisted of 8% (w/v) PEG (MW 35 kDa) and 17% (v/v) Optiprep densifier dissolved in mTeSR medium. For cell encapsulation, flow rates of the core, shell, shielding oil, and crosslinker oil streams were maintained at 4, 4, 50, 60 μL/min, respectively. The microcapsules were generated at the frequency of ~10 capsules/s and typically had ~200 cells per capsule. A core solution containing cell suspension first passed through the filter device to trap cell clumps and was then injected into the encapsulation device (see Fig. S2B for description of the filter device). The microcapsules carrying HUES-8 cells were collected into 15 mL tube filled with 5 mL E8 medium (STEMCELL Technologies), then distributed into a 6 well-plate at density of 2000 capsules/well and incubated at 37 □ with 5% CO_2_. The culture medium was changed every 24 h. The same encapsulation process was followed for Hep and PEG microcapsules. As a control condition, HUES-8 cells were seeded in Aggrewell (AggreWell™400; STEMCELL Technologies) at a density of ~200 cells/well. Protocols used for cultivation of encapsulated and unencapsulated HUES-8 spheroids are described in the following sections.

### 2.7. Viability and proliferation of encapsulated HUES-8

Stem cell viability inside the capsules was analyzed by a Live/Dead assay (Thermo Fisher Scientific), where live and dead cells fluoresce in green and red, respectively. In brief, the capsules with encapsulated spheroids were rinsed with PBS then incubated in 1x PBS containing 4 μM calcein-AM and 2 μM ethidium homodimer-1 (EtBr-1) for 30 min. After washing with 1x PBS, images of the encapsulated spheroids were obtained using an inverted fluorescence microscope. Cell viability was quantified by calculating the ratio of the live cells over the total number of cells in the images. The size of the HUES-8 spheroids in the microcapsules was evaluated at 1, 3, 5, and 7 days of culture with an ImageJ software (n=20 capsules per time point) from the images.

### 2.8. Pluripotency maintenance of encapsulated HUES8 cells

To test the effect of one-time loading of pluripotency signals, microcapsules carrying HUES-8 cells were immersed in E8 medium for 1 h immediately after encapsulation and then transferred into E6 medium (without FGF-2 and TGF-β1, STEMCELL Technologies) for subsequent cultivation (2000 microcapsules per well). E6 medium was exchanged every 24 h. This one-time loading or immobilization was performed with microcapsules with Hep and PEG, and conditions that use these capsules were termed as Hep^Imm^ and PEG^Imm^ conditions, respectively. Additional experimental groups included HUES-8 spheroids in 3D Aggrewell plates, and PEG capsules and Hep capsules exposed to soluble FGF-2 and TGF-β1 in E8 medium over the course of 5 days. These experimental groups were termed as bare spheroid, PEG^Sol^ and Hep^Sol^ respectively. When exchanging medium, the microcapsules were collected in a 100 μm cell strainer to remove used medium and washed with fresh medium twice. Then, 2000 microcapsules were placed into a 6-well plate containing 2 mL of fresh medium.

### 2.9. Differentiation of encapsulated HUES8 cells into definitive endoderm

A standard endodermal differentiation protocol calls for HUES-8 spheroids to be cultured in MCDB131 medium (Thermo Fisher Scientific) containing 200 ng/mL Nodal or 100 ng/mL Activin A (both obtained from R&D Systems) and 3 μM CHIR99021 (Stemgent, Cambridge, MA, USA) [12, 29]. Because Activin A does not possess heparin-binding domains, it was replaced by Nodal for in-capsule differentiation. Nodal is a ligand very similar to Activin A that has been shown to signal through the same receptors and has been used to induce endodermal differentiation [30]. To test effects of one-time loading of this inductive cue, PEG or Hep microcapsules carrying HUES-8 spheroids were placed into 200 ng/mL Nodal in MCDB131 medium for 1 h. Subsequently, microcapsules were transferred into a 6-well plate containing MCDB131 medium supplemented with CHIR99021 but without Nodal. Experimental groups testing the effects of one-time loading of Nodal were as follows: Hep^Sol/Imm^ and Hep^Imm/Imm^. Additional experimental groups were designed to test exposure to soluble endodermal signal: PEG^Sol/Sol^ and bare spheroids. In this nomenclature, the abbreviations before and after the forward slash refer to the mode of exposure to pluripotency and endodermal signals, respectively. For example, Hep^Sol/Imm^ condition denotes exposure to soluble pluripotency signals followed by immobilization of endodermal signal Nodal, whereas Hep^Imm/Imm^ means that pluripotency and endodermal signals were immobilized into microcapsules. We also note that bare HUES-8 spheroids maintained in 3D culture plates (Aggrewell) were exposed to soluble Nodal in MCDB131. All endodermal differentiation experiments lasted for 3 days.

### 2.10. RT-PCR analysis of pluripotency and endodermal gene expression

To examine the gene expression levels of HUES-8 cells, 500 microcapsules were first broken down by applying electronic pestle for 3 min, then total RNA was extracted using a commercial kit (Qiagen, Valencia, CA, USA) following manufacturer’s instructions. Approximately 100 ng of total RNA was processed using reverse transcription kit to synthesize cDNA (Roche, Basel, Switzerland). The primer sequences used for RT-PCR analysis are listed in Table S1. Gene expression analysis was performed with the QuantStudio™ 5 System (Thermo Fisher Scientific) using SYBR Green (Roche) and was normalized to glyceraldehyde 3-phosphate dehydrogenase (GAPDH). Real-time PCR was performed with an amplification procedure consisting of 40 cycles of denaturation at 95 □ for 5 s, annealing at 55 □ for 15 s, and extension at 69 □ for 20 s. The final analysis was performed based on the threshold cycles using the ΔΔC_T_ method.

### 2.11. Immunofluorescence staining of hPSC spheroids

The encapsulated spheroids were fixed with 4% paraformaldehyde and immersed in 30% sucrose in PBS for 24 h. The spheroids were then imbedded in OCT compound (Fisher Healthcare, Waltham, MA, USA) and sectioned into 10 μm thick slices using a cryostat instrument (Leica CM1950; Leica Biosystems Inc., Buffalo Grove, IL, USA). Then the sections were permeabilized by immersion for 20 min in 1x PBS supplemented with 0.1% Triton X-100 in 2% BSA, and blocked with 2% BSA in 1x PBS for 2 h at 25°C. After thorough washing with 1x PBS, the slides were incubated in 5 μg/mL solution of anti-SOX2 antibodies (R&D Systems) or 5 μg/mL solution of anti-Sox17 (SantaCruz, Dallas, TX, USA) for 1 h at 25°C. Both antibody solutions were prepared in a blocking buffer of 2% BSA in 1x PPBS. The sections were washed again three times for 5 min each, then incubated with the corresponding secondary antibodies conjugated to Alexa Fluor 488 for Sox2 and 694 for Sox17 (2 μg/mL; Invitrogen) for 1 h at 25°C in the dark. A sectioned slice was immersed in 50 μL of mounting medium containing DAPI (Vectorlab, San Francisco, CA, USA) placed under a coverslip (170 μm from Fisher Healthcare) and imaged using an Olympus fluorescence microscope (see above for microscope information).

### 2.12. Modeling growth factor uptake and release

To evaluate loading and release of FGF-2 and TGF-β1 in a bioactive capsule, we used COMSOL Multiphysics^®^ software (Version 5.6, Burlington, MA). We modeled the loading and release using diffusion (medium outside the capsule, shell, and core) and reaction (shell) processes. One-hour loading was implemented by using initial concentrations of the GF in the medium outside the capsule (6.06 and 0.156 nM of FGF-2 and TGF-β1, respectively) that were reset to 0 after 1 hour. The release was followed for a period of 12 hours. The reaction kinetics for GF binding with heparin were modeled using a reversible kinetic expression: 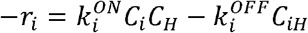. Here, *C_i_, C_H_, and C_iH_* are concentration of the GF, heparin, and GF-heparin complex. 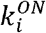 and 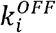 are rate constants [31]. Since the concentrations of GF were in the nM range, to improve accuracy of the solver, the reaction diffusion equations were non-dimensionalized prior to implementation in COMSOL. Microcapsules in a confined medium were simulated using experimental parameters of 3600 microcapsules (400 μm in diameter) distributed in 2 mL of medium. This led to a fluid layer of about 1.0 mm diameter surrounding each capsule with a no flux boundary condition at the imaginary interface between adjoining fluid layers. Cellular spheroid was simulated as an inert sphere of 300 μm diameter concentrically positioned inside the microcapsule. The reaction-diffusion equations for GF, heparin, and GF-heparin complex were solved using 2D axisymmetric model in COMSOL. To avoid instability, a small diffusivity of 10^-15^ m^2^/sec was used for heparin and GF-heparin complex in the medium and shell. Each GF loading and release were simulated separately. Table S2 lists all the parameters used in the simulations.

### 2.13. Statistical analysis

All experiments were performed four times or more. Data were statistically evaluated by Student’s t-test. All data were presented as mean ± standard deviation of the mean. The minimum level of significance was set at p < 0.05 (*: p < 0.05, **: p < 0.01, and ***: p < 0.001).

## 3. Results and Discussion

The goal of this paper was to develop bioactive core-shell microcapsules to 1) promote formation of hPSC spheroids, 2) achieve sequential loading of GFs to mimic multi-step maintenance and differentiation protocols (Scheme 1), and 3) enable local and sustained release of GFs to the encapsulated hPSCs.

### 3.1 Fabricating bioactive core-shell microcapsules

Heparin was functionalized by transesterification reaction with methacrylic anhydride (Fig. S3A) to synthesize bioactive and crosslinkable Hep-MA moieties. The degree of methacrylation (~30%) was established with ^1^H-NMR spectroscopy by comparing methacrylate peaks of MA at ~5.6 and ~6.1 ppm and proton peaks on the repeating disaccharide units of heparin before and after functionalization (Fig. S3B) [32].

Hep-MA molecules were used in the microfluidic capsule fabrication process described in Fig. 1A. A co-axial flow focusing microfluidic device was used to fabricate microcapsules with core-shell architecture. Two aqueous streams were injected into the microfluidic device: core solution carrying stem cell suspension and shell solution containing PEG4MAL and Hep-MA. The core solution comprised of non-reactive PEG (MW 35 kDa) and Optiprep densifier, and had viscosity of 20 mPa·s, similar to that of the shell stream. As shown in the cross-section view in Fig. 1A, channels for core and shell flow streams were 120 μm and 200 μm in height, respectively. Upon entering the 300 μm-tall oil channel, aqueous co-axial flow streams became discretized into droplets with a shell region wrapping around the core. Two oil junctions were used in this device (see Fig. 1A and 2A); the first oil junction served to discretize, shield, and stabilize core-shell droplets, and the second oil junction was used to deliver a di-thiol crosslinker, DTT, that reacted with PEG4MAL and Hep-MA *via* click and Michael addition reactions, respectively. This resulted in formation of a thin (~15 μm) PEG hydrogel network with incorporated heparin moieties (Fig. 1B). An encapsulation device and the process of encapsulation are shown in Fig. 2A and Supporting Video 1, respectively.

**Fig. 1.**
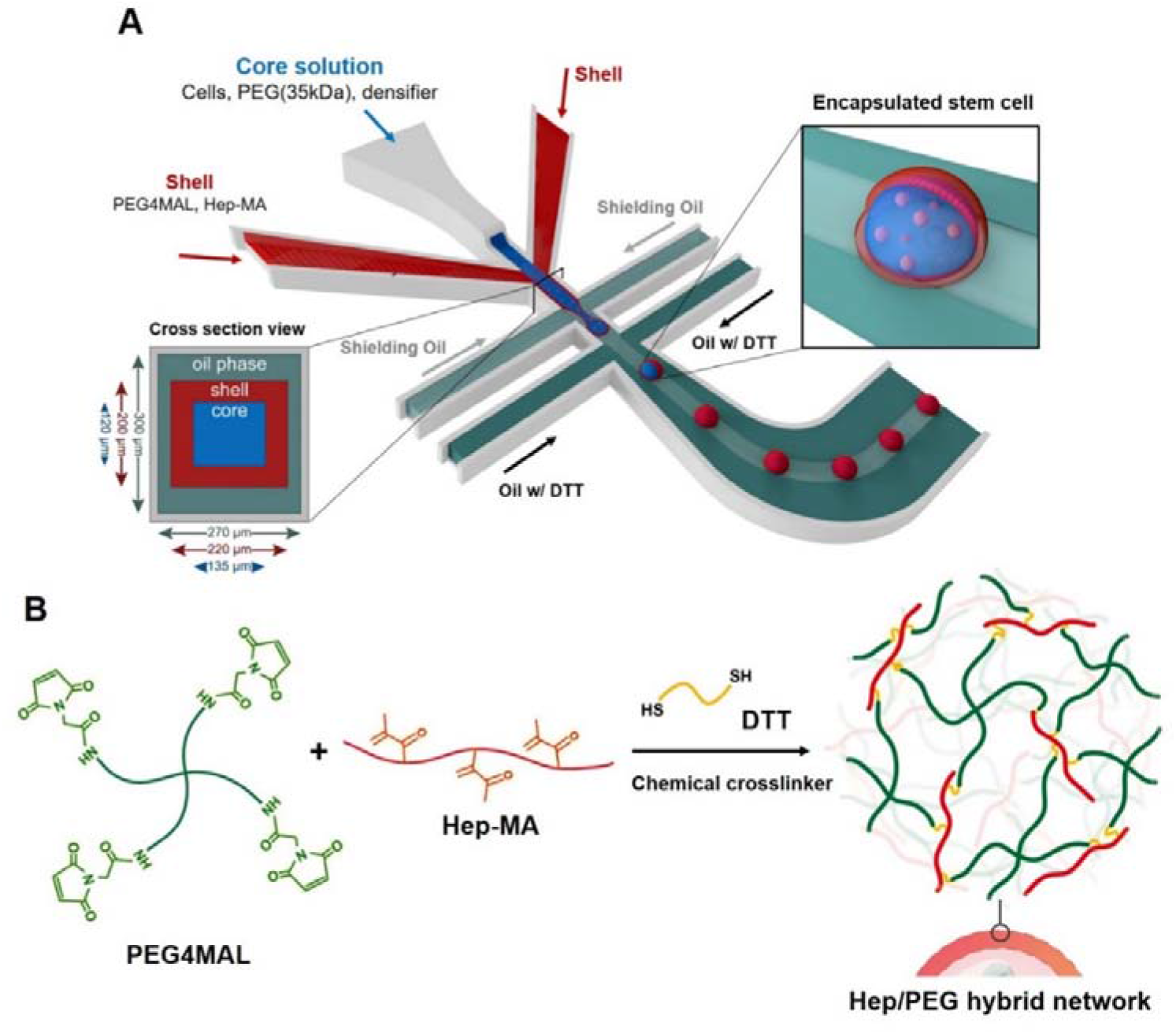
Fabrication of bioactive microcapsules. (**A**) Microfluidic flow-focusing device is infused with two aqueous streams: one containing stem cell suspension, and another carrying methacrylated heparin (Hep-MA) and 4-arm PEG-maleimide (PEG4MAL) mixture. Emulsification occurs by exposure to two oil streams; the first designed to stabilize aqueous droplets and the second to deliver a crosslinker DTT to Hep-MA and PEG4MAL. The core, shell, and oil channels have heights of 120 μm, 200 μm, and 300 μm, respectively. (**B**) Chemical crosslink mechanism between shell polymers (Hep-MA and PEG4MAL) and dithiothreitol (DTT) as crosslinker in oil.

**Fig. 2.**
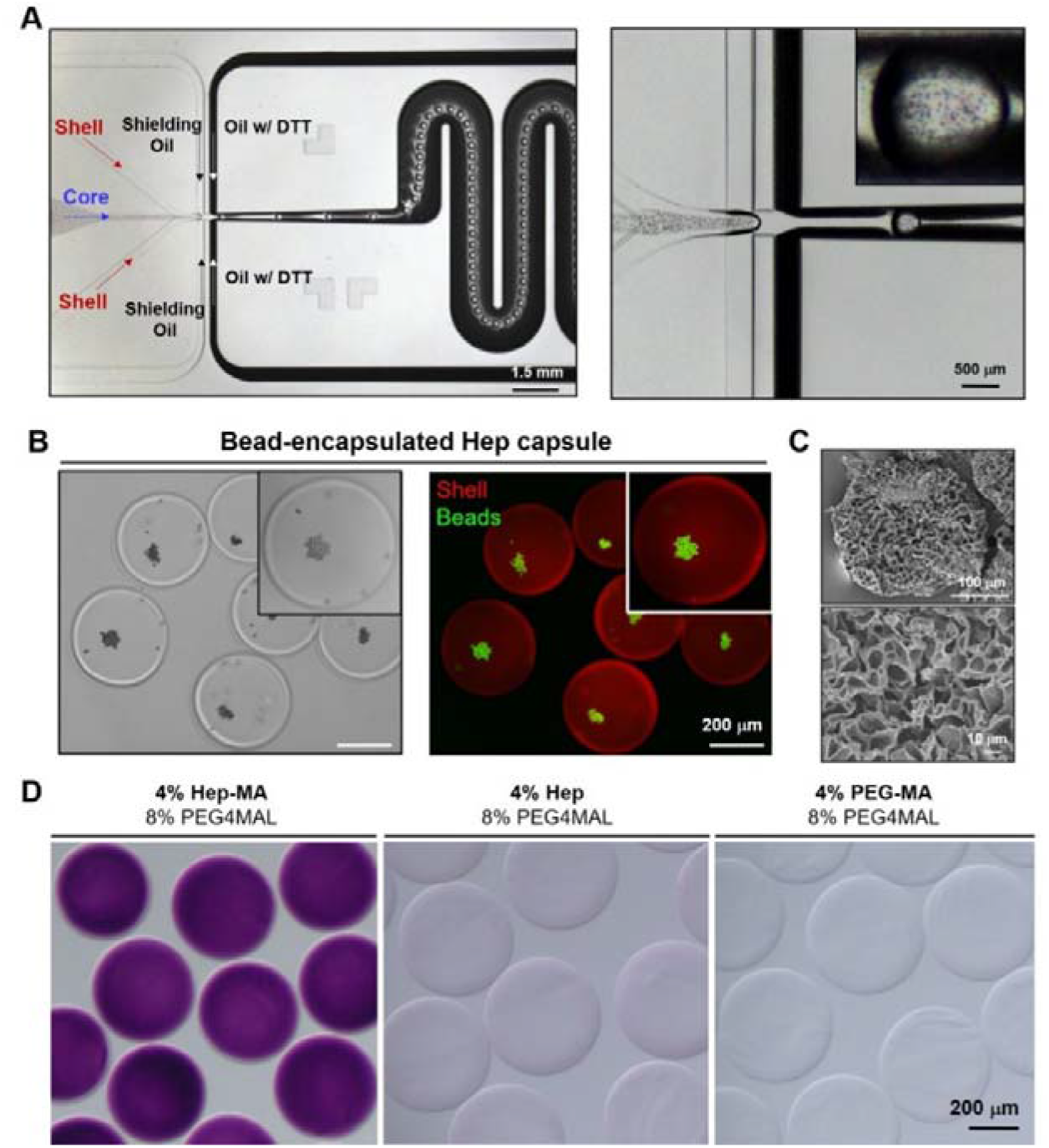
Characterization of core-shell microcapsules. **(A)** Top view of coaxial flow-focusing device during the encapsulation of microbead in core-shell capsules. **(B)** The encapsulated microbeads (green) are accumulated in the liquid core region while the shell region is clearly distinguished in red. **(C)** SEM images of the freeze-dried core-shell microcapsule. **(D)** Heparin-based core-shell microcapsules stained with positively charged toluidine blue O. Core-shell microcapsules with physically incorporated heparin or no heparin did not retain toluidine blue O dyes.

Non-reactive high viscosity molecules leached out from the core and were replaced by water molecules after the microcapsules were transferred into aqueous medium. This led to the formation of microcapsules with a thin hydrogel shell and an aqueous core [12, 15]. We should note that while microfluidic channels were designed to fabricate capsules with 300 μm diameter, the final capsule dimension reached 380 μm diameter upon swelling in the aqueous environment.

### 3.2 Characterization of core-shell architecture and heparin incorporation into microcapsules

In order to confirm core-shell structure of microcapsules, fluorescein-labeled microbeads and rhodamine-PEG-SH were included in the core and shell streams of the microfluidic device. Fig. 2B shows that with microcapsules resting at the bottom of a Petri dish, microbeads aggregated in the center of each capsule. This suggested that microbeads were free to move and aggregate, and thus were surrounded by an aqueous environment inside the microcapsules. Conversely, microbeads remained dispersed and immobile when encapsulated into microcapsules with hydrogel core (data not shown). While aggregation of microbeads confirmed an aqueous core, immobilization of rhodamine-PEG suggested the presence of a thin (~15 μm) hydrogel shell (see Fig. 2B). It is worth noting that shell thickness could be varied from 15.5 ± 1.5 to 40.0 ± 2.5 μm, and core diameter for 350.0 ± 8.5 to 255.0 ± 6.5 μm by adjusting the core and shell flow rates (Fig. S4). Because cells tended to become entrapped in thicker shells approaching 40 μm (data not shown), we chose to operate microfluidic encapsulation device with both core and shell streams at the same flow rate (4 μL/min). These operational conditions resulted in microcapsules with ~15 μm hydrogel shell and ~380 ± 12 μm in diameter. The surface topography of the core-shell microcapsules was characterized by SEM (Fig. 2C). The surface of the lyophilized microcapsules was porous with pores ranging from 10 to 30 μm. This structure was similar to that of other lyophilized hydrogels [33].

We characterized diffusion of dextrans of different molecular weights from Hep capsules. Results presented in Fig. S5 and Table S3 demonstrate that fluorescent dextran molecules of similar MW to GFs of interest were released within tens of minutes of loading. These results indicate that Hep hydrogel capsules were porous and that dextran molecules comparable to GFs in size, but not in biomolecular composition, diffused out rapidly. These observations underscore the fact that sustained release of GFs described in later sections of this study is governed by binding to heparin moieties in microcapsules and not by soluble GFs trapped within the microcapsule.

Presence of negatively charged heparin moieties was confirmed by staining with positively charged dye toluidine blue O (see Fig. 2D). Incorporation of Hep-MA into the shell flow stream resulted in microcapsules that were stained strongly with toluidine blue O (purple color). Conversely, inclusion of heparin without methacrylate groups into the shell stream resulted in microcapsules that showed no toluidine blue O staining, suggesting that unfunctionalized heparin leached out from the microcapsules (see Fig. 2D). Assuming that concentration of Hep in the gel is similar to solution concentration and taking into account 15 μm thickness of the shell, we estimate 1.071 x 10^13^ heparin molecules per capsule.

The number of heparin molecules per microcapsule is a function of both the thickness of hydrogel shell and heparin content in the shell stream carrying prepolymer molecules. As noted above, we determined empirically that shells with thickness exceeding 15 μm contained embedded cells that did not participate in the aggregation within the core. We deemed it important to maximize aggregation of cells in the microcapsules and proceeded to use shell thickness of 15 μm for subsequent experiments. In terms of varying heparin content, we attempted to fabricate microcapsules with 2%, 4% and 6% w/v heparin while keeping 8% w/v PEG4MAL constant. We determined that capsules with the highest heparin content (6%) were not mechanically stable (data not shown), while 2% and 4% heparin prepolymer produced mechanically robust hydrogel capsules. Based on these empirical observations, we chose the prepolymer solution containing 4% Hep-MA and 8% PEG4MAL for fabricating microcapsules in this study.

### 3.3 Growth factor release from bioactive microcapsules

After demonstrating incorporation of heparin moieties into core-shell microcapsules, we proceeded to characterize loading and release of GFs from these bioactive capsules. In this study, we focused on GFs that induce pluripotency, FGF-2 and TGF-β1 [34–36], as well as on endodermal signal, Nodal [14, 37]. All three GFs contain heparin-binding domains [31]. Release profiles of GFs loaded into Hep microcapsules were compared to PEG microcapsules. As may be appreciated from Fig. 3A, 90% of GF molecules loaded into PEG microcapsules were detected after the first 30 min, indicating burst release. The release from PEG microcapsules was indicative of diffusion and was consistent with dextran diffusion studies described in Fig. S5. In contrast, continued and sustained release was observed from Hep microcapsules suggesting that affinity interactions with heparin moieties governed GF release from bioactive microcapsules.

**Fig. 3.**
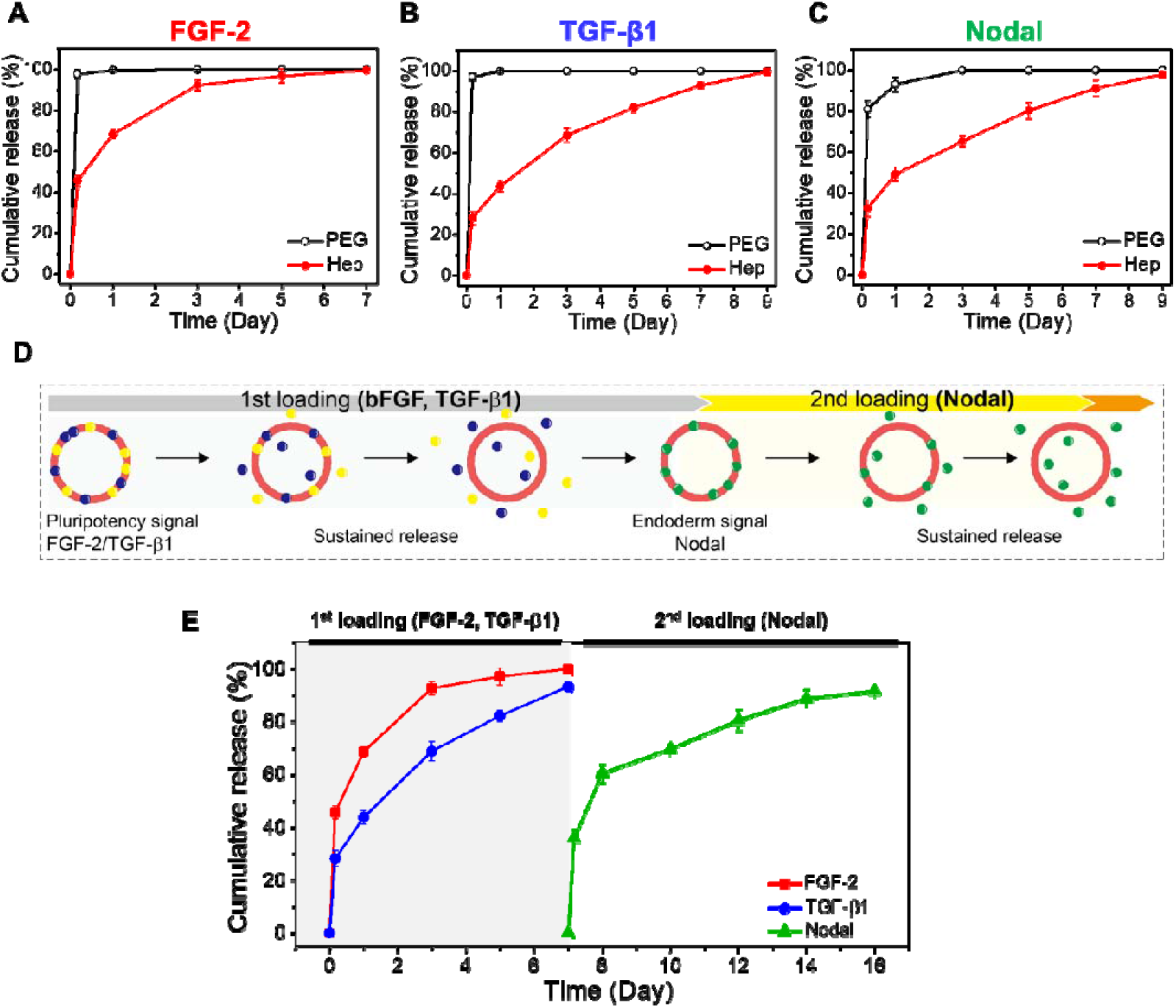
Growth factor release from bioactive microcapsules. Release profile of (**A**) FGF-2, (**B**) TGF-β1, and (**C**) Nodal from Heparin capsules (Hep) and inert PEG capsules (PEG). (**D**) Workflow for the release and reloading of different GFs into heparin capsules at different stages. (**E**) Sequential release profile from subsequent reloading: 1^st^ loading (FGF-2 & TGF-β1), and 2^nd^ loading (Nodal) of Hep capsules.

GF concentration and incubation time were important parameters to consider for loading and release experiments. For the sake of simplicity and consistency, GF concentrations for loading into microcapsules were identical to those present in solution for standard maintenance and differentiation protocols. We tested multiple loading times ranging from 1 h to 24 h and determined that 90% of maximal FGF-2 loading occurred within 1 h of incubation (see Fig. S6) and proceeded to use 1 h loading time throughout this study. As highlighted by the GF loading results (see Table 1), bioactive Hep microcapsules retained 3 to 5 times the amount of GF present in PEG microcapsules. These data once again highlight the bioactive nature of Hep microcapsules and their enhanced capacity for sequestering GFs.

**Table 1.** Growth factor loading into microcapsules. After incubation in GF-containing buffer, microcapsules were removed and the amount of GF remaining in solution was quantified by ELISA. Amounts of GF are reported per 3600 microcapsules.

| GF type | Soluble GF amount (ng) | Loaded GF amount PEG capsule (ng) | Loaded GF amount Hep capsule (ng) | Hep capsule: Nodal loaded after release of FGF-2 and TGF-β1 (ng) |
| --- | --- | --- | --- | --- |
| FGF-2 | 100 | 15 | 47 | NA |
| TGF-β1 | 2 | 0.3 | 1.5 | NA |
| Nodal | 200 | 38 | 133 | 125 |

Beyond enhanced loading, bioactive microcapsules enabled controlled release of GFs over the course of 7 to 9 days. 90% release was observed at day 7 for FGF-2 (see Fig. 3A), day 9 for TGF-β1 (see Fig. 3B), and day 9 for Nodal (see Fig. 3C). These multi-day release profiles were in stark contrast with rapid (within 30 min) release of GFs from biologically inert PEG microcapsules.

We should note that the key advantage of Hep microcapsules is reversibility of heparin-GF interactions which means that capsules vacated by one type or set of GFs may be loaded with another set of GFs to mimic multi-step stem cell cultivation protocols. Fig. 3(D, E) describe an experiment that tested the possibility of loading, releasing, and reloading different types of GFs into the same microcapsules. As shown in Fig. 3D, Hep microcapsules were first loaded with pluripotency inducing signals, FGF-2 and TGF-β1, and then, 7 days later, were reloaded with endodermal signal, Nodal. As seen from Fig. 3E, this two-step loading and release process produced profiles similar to those observed for GFs loaded individually into pristine microcapsules (see Fig. (3A, B, C)). Incubation of Hep microcapsules with FGF-2 or TGF-β1 individually or as a mixture resulted in similar GF loading (see Table 1). Even more interestingly, the same amount of Nodal was incorporated into pristine Hep microcapsules and microcapsules that previously contained FGF-2 and TGF-β1. This result highlighted that Hep microcapsules remained bioactive and capable of sequestering Nodal after 9 days of storage at 37°C in PBS.

### 3.4 Use of bioactive microcapsules for cultivation of hPSCs

After characterizing GF loading and release from Hep microcapsules, we proceeded to assess the utility of these bioactive microcapsules for maintenance and differentiation of hPSCs, using hESC line (HUES-8 cells) as a model. We previously described a microfluidic encapsulation system that included a filter device located upstream of the flow-focusing encapsulation device [12]. The filter device was further refined in the present study. It was comprised of a flow channel with an array of triangle-shaped posts measuring 200 μm per side, and with pitch ranging from 400 μm at the inlet to 30 μm at the outlet (see Fig. S2). COMSOL modeling was used to estimate the shear stress to be 0.08 and 2.77 dyn/cm^2^ at the inlet and outlet of the filter device, respectively, for the flow rate of 4 μL/min (see Fig. S2). The highest shear stress created in the filter device is well below the threshold of 30 dyn/cm^2^ above which cell damaged has been reported [38]. The use of the filter device allowed us to improve uniformity of cell loading into capsules (cell number per capsule) by trapping cell aggregates, and increase cell occupancy of capsules (see ref [12] for comparison of cell encapsulation with and without the filter device). In the present study, the encapsulation system comprised of the two microfluidic devices allowed us to achieve 90% capsule occupancy in a typical encapsulation run, with spheroid diameter being 102.5 ± 9.5 μm. Importantly, the process of filtering and encapsulation did not compromise viability of HUES-8 cells. Live/dead staining revealed that encapsulated HUES-8 cells had viability of 94 ± 3.1% (see Fig. 4A), which was comparable to the viability of cells before encapsulation (94.5 ± 1.4%, see Fig. 4B).

**Fig. 4.**
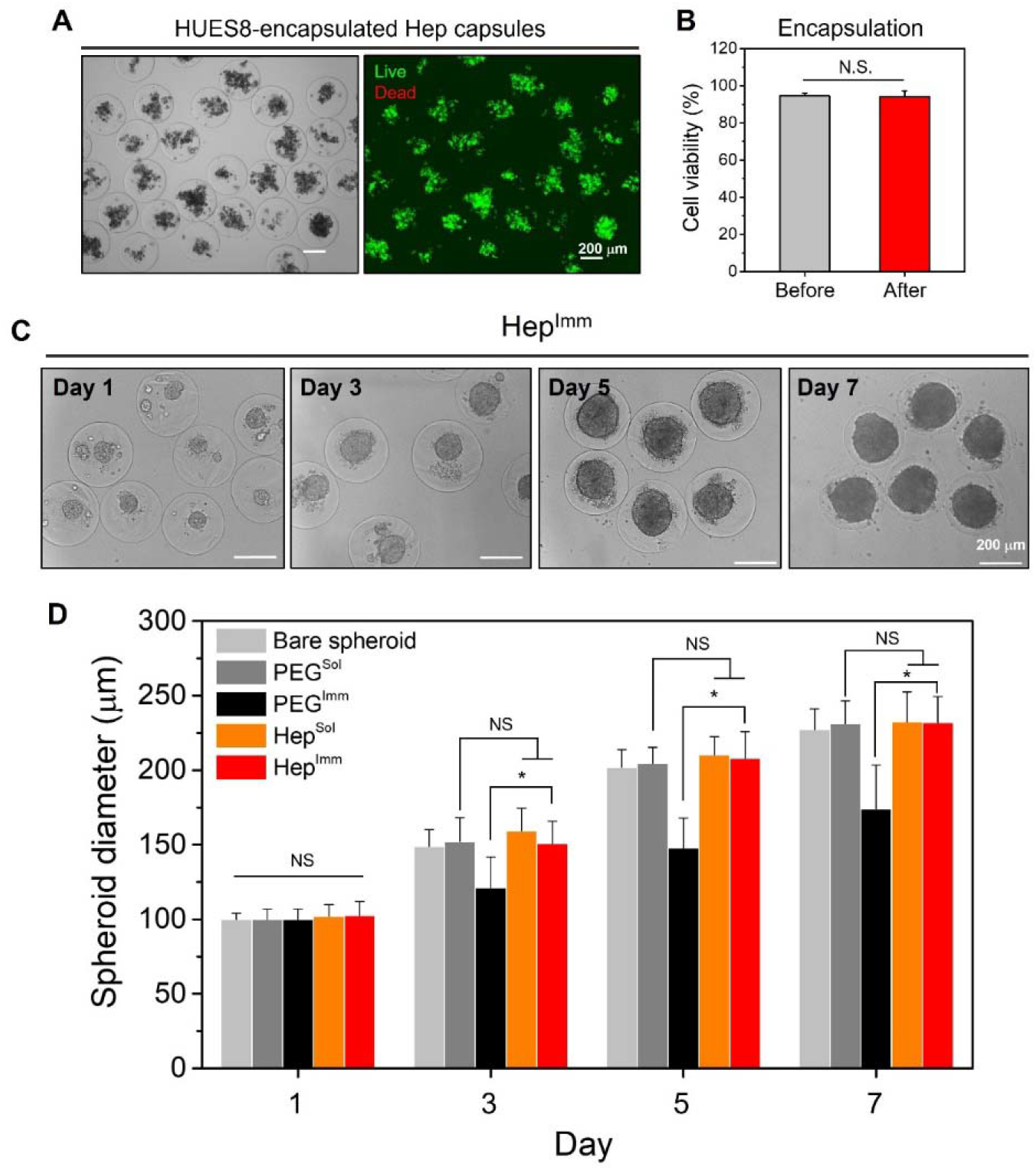
Encapsulation and growth of hPSC spheroids. (**A**) Brightfield image and live/dead fluorescent image of encapsulated stem cells. (**B**) Comparison of viability before and after encapsulation of HUES-8 (n=30 capsules, p < 0.05). (**C**) Representative brightfield images at different time points during cultures. hPSC spheroids were maintained in pluripotency medium. (**D**) Size distribution of encapsulated hPSC spheroids and bare spheroids at different time points during cultures. Sol refers to HUES-8 exposed persistently to soluble FGF-2 and TGF-β1 with daily medium exchange. Imm refers to HUES-8 cells exposed momentarily to GFs for GF loading within the microcapsules. Bare spheroids (without capsules) were cultured in commercial 3D plates (Aggrewell) and exposed to soluble GFs. Data are presented as mean ± standard deviation and statistically evaluated by Student’s t-test. Each group was compared to Hep^Imm^ with the minimum level of significance set at p < 0.05 *.

Having verified viability of encapsulated HUES-8 cells, we proceeded to characterize effects of GF incorporation on stem cell proliferation and phenotype expression. Our experimental groups consisted of a bare hPSC spheroid control group cultured in a commercial 3D plate (Aggrewell) and encapsulated groups that were further subdivided based on the bioactivity into Hep vs. PEG microcapsules and based on the mode of GF exposure into Sol or Imm groups.

As the first step in stem cell characterization, we compared spheroid formation and proliferation for these groups (see Fig. 4D and S7). Importantly, the experiment was designed to ensure that the bare spheroids formed in commercial 3D plates were of similar size to stem cell spheroids inside capsules. In the case of bare spheroids as well as PEG^Sol^ and Hep^Sol^ microcapsules, HUES-8 spheroids were cultured in pluripotency maintenance medium (E8) containing FGF-2 and TGF-β1 for the duration of the 7-day experiment. For PEG^Imm^ and Hep^Imm^ conditions, encapsulated stem cell spheroids were exposed to E8 medium for 1 h and then switched to E6 media lacking FGF-2 and TGF-β1. Spheroid formation and growth associated with these 5 experimental groups were characterized and quantified by microscopy. As seen from Fig. 4(C, D) and Fig. S7, spheroid formation and growth were similar in all experimental groups except for PEG^Imm^ condition where hPSCs were exposed to pluripotency signals for short duration (1 h). While hPSC spheroids in Hep^Imm^ condition experienced similarly short exposure to FGF-2 and TGF-β1, these spheroids grew over the course of the subsequent 6 days at a rate comparable to that of spheroids exposed to soluble GFs (see Fig. 4D). We note that when cells were fed with soluble GFs, the media was changed every 24 h. This result suggested that one-time incorporation of pluripotency signals into Hep microcapsules produced a lasting effect on hPSC growth over the course of 7 days. It is also worth noting that the observation of spheroid growth over time aligned well with GF release which also occurs on the timescale of 7 to 9 days (See Fig. 3E). In the experiments described below, we assess the pluripotency state of these encapsulated hPSCs.

### 3.5 Assessing pluripotency of encapsulated hPSCs

We relied on the combination of gene expression analysis by RT-PCR and immunofluorescence staining to assess pluripotency of hPSCs. As described in the process flow diagram (Fig. 5A), hPSCs spheroids were maintained under conditions inducing pluripotency for 5 days. Having noted that spheroid formation and proliferation was least effective for PEG^Imm^ microcapsules, we eliminated this experimental group from pluripotency and differentiation studies. Thus, we tested four experimental groups: 1) bare spheroids, 2) PEG^Sol^ and 3) Hep^Sol^ microcapsules exposed to E8 medium (with FGF-2 and TGF-β1) for the duration of the experiment with daily medium exchange, and 4) Hep^Imm^ capsules loaded with FGF-2 and TGF-β1 for 1 h followed by cultivation in E6 medium without inductive GFs. RT-PCR analysis for markers of pluripotency (Sox2, Oct4, and Nanog) revealed significant differences between the four experimental groups analyzed (see Fig. 5B). Hep microcapsules with immobilized GFs (Hep^Imm^) induced high levels of pluripotency gene expression similar to those observed for Hep^Sol^ or PEG^Sol^ microcapsules, and higher than those of bare spheroids. While it may have been expected that PEG^sol^ and bare spheroid conditions elicit similar levels of pluripotency gene expression, there were notable differences between these experimental groups. While encapsulated stem cell spheroids were placed into filter inserts and then moved from well to well during media exchange, bare spheroids resided in pyramidal wells of a commercial 3D culture plate where media exchange could not be accomplished as efficiently. We estimate that 50 to 75% of media was exchanged for bare spheroid experimental group.

**Fig. 5.**
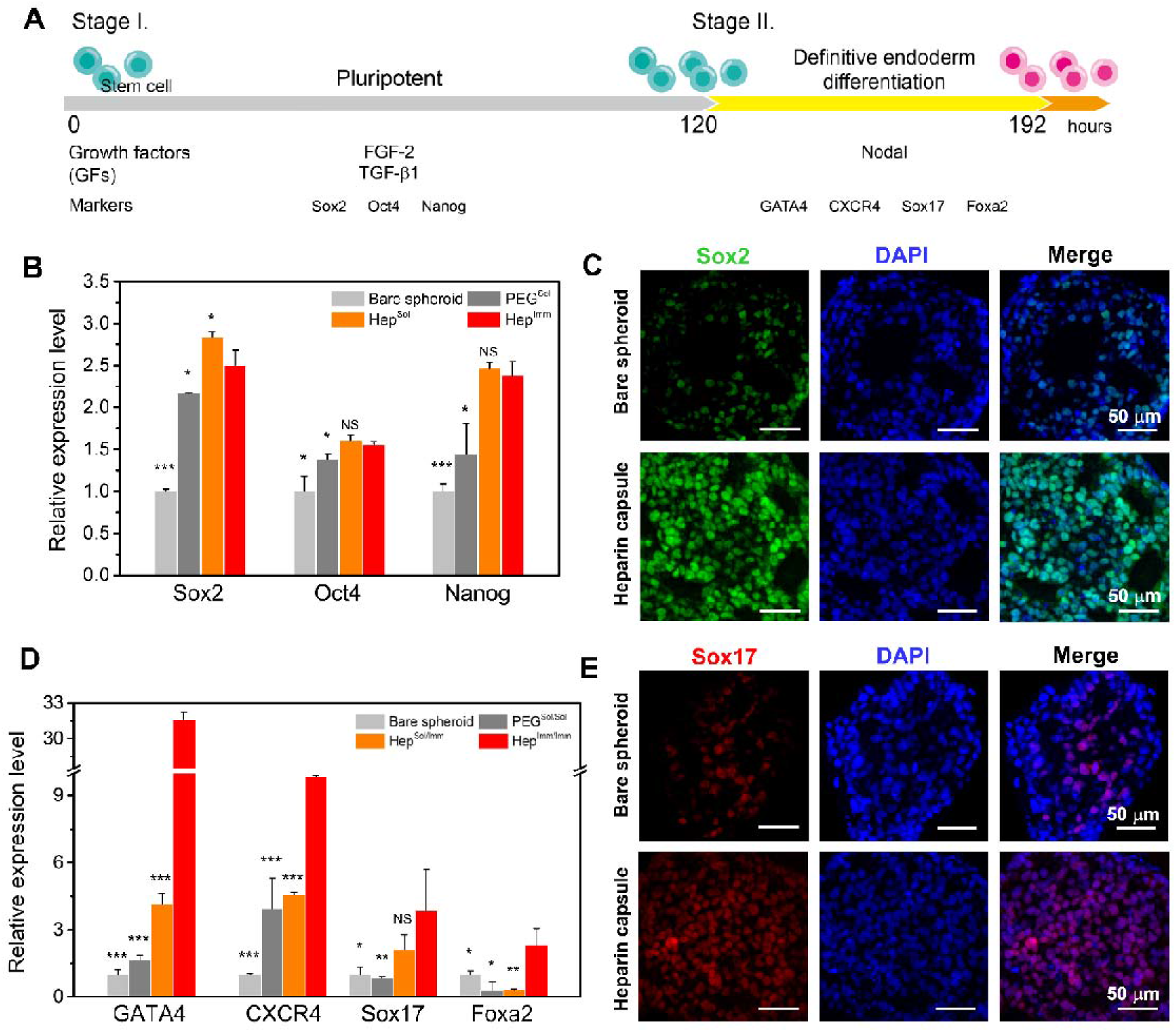
Pluripotency and definitive endoderm expression in bioactive microcapsules. (**A**) Timeline for two-step definitive endoderm (DE) differentiation protocol of encapsulated HUES-8 cells. (**B**) Pluripotency gene expression (Sox2, Oct4, and Nanog) for encapsulated and bare HUES-8 spheroids. Sol refers to HUES-8 exposed persistently to soluble FGF-2 and TGF-β1 with daily medium exchange. Imm refers to HUES-8 cells exposed momentarily to GFs for GF loading within the microcapsules. Bare spheroids were cultured in Aggrewell without encapsulation and exposed to soluble GFs persistently. (**C**) Immunofluorescent staining for Sox2 expression of bare spheroids (top) and encapsulated spheroids in heparin microcapsules (bottom). (**D**) Expression of definite endoderm markers (GATA4, CXCR4, Sox17, and Foxa2) for 3D encapsulated and bare HUES-8 spheroids exposed to Nodal. (**E**) Immunofluorescent staining for Sox17 expression of bare spheroids (top) and encapsulated spheroids in heparin microcapsules (bottom). Data of (B) and (D) are presented as mean ± standard deviation and statistically evaluated by Student’s t-test. Each group was compared to Hep^Imm^ (B) and Hep^Imm/Imm^ (D) with the minimum level of significance set at p < 0.05 *: p < 0.05, **: p < 0.01, and ***: p < 0.001.

Immunofluorescence staining for Sox2 was used to confirm gene expression analysis. As seen from Fig. 5C, stronger Sox2 staining was observed for stem cell spheroids in Hep^Imm^ capsules compared to bare spheroids, confirming higher level of pluripotency in the former condition.

In summary, our results demonstrate that one-time loading followed by sustained, local release of inductive cues was sufficient to maintain pluripotency of the encapsulated HUES-8 spheroids over the course of 5 days. The reasons for high level of pluripotency maintenance in bioactive microcapsules are likely two-fold: 1) high local concentration of inductive cues and 2) improved potency/bioavailability of GFs. For example, while free FGF-2 has a relatively short half-life of 85 min due to proteolytic degradation and denaturation, heparin-bound FGF-2 has been shown to be more stable [39, 40].

### 3.6 Endodermal differentiation of hPSC spheroids in bioactive microcapsules

While pluripotent stem cells may be differentiated into any adult cell type, our labs are interested in pancreatic and hepatic lineages – both of which originate from the endoderm [12, 29, 41]. Therefore, we evaluated endodermal differentiation of hPSC spheroids in bioactive microcapsules. Typical differentiation protocols rely on Activin A, a member of TGF-β superfamily, for driving endodermal differentiation [42, 43]. However, Activin A does not possess heparin-binding domains [44] and may not be loaded into bioactive microcapsules. Therefore, in this study, we used Nodal, which is closely related to Activin A [30] and has also been used for endodermal differentiation of hPSCs [37]. Nodal does possess heparin-binding domains and has been shown by us to have affinity for heparin-modified surfaces and heparin-based hydrogels [14].

From a technical standpoint, we wanted to demonstrate that bioactive microcapsules may be used to mimic traditional differentiation protocols where inductive cues are introduced sequentially for prescribed periods of time. To prove the concept, in this study we pursued a two-step in-capsule cultivation protocol where loading and release of pluripotency cues (FGF-2 and TGF-β1) was followed by loading and release of endodermal cue, Nodal (see Fig. 5A). As before, we compared four experimental groups, however, our nomenclature evolved to account for two sets of GFs and Imm vs. Sol exposure. The experimental groups analyzed for endodermal differentiation were 1) bare spheroids, 2) PEG^Sol/Sol^ microcapsules exposed to soluble pluripotency and endodermal signals, 3) Hep^Sol/Imm^ where exposure to soluble pluripotency cues was followed by incorporation of Nodal, and 4) Hep^Imm/Imm^ where one-time loading of pluripotency cues was followed by one-time loading of Nodal. RT-PCR analysis (see Fig. 5D) revealed that the best endodermal gene expression was observed in Hep^Imm/Imm^ microcapsules, followed by Hep^Sol/Imm^ capsules, PEG^Sol/Sol^ capsules, and bare spheroids. These results indicated that loading of Nodal into bioactive Hep microcapsules led to better endodermal differentiation compared to continuous exposure of hPSCs to soluble Nodal either in PEG microcapsules or in bare spheroids. The differences between Hep^Sol/Imm^ and Hep^Imm/Imm^ capsules are intriguing and warrant further discussion. Hep^Sol/Imm^ microcapsules were exposed to fresh E8 medium containing soluble pluripotency signals, FGF-2 and TGF-β1, daily during pluripotency maintenance. This means that Hep^Sol/Imm^ microcapsules retained a high concentration of pluripotency GFs when they were transferred from E8 medium into endoderm medium. Therefore, delivery of pluripotency signals to the encapsulated hPSCs may have continued during differentiation and may have contributed to lower efficiency of endoderm expression. An alternative explanation may be that Nodal molecules were not loaded effectively in Hep^Sol/Imm^ microcapsules because heparin sites were occupied by FGF-2 and TGF-β1 molecules. This underscores the importance of timing for loading GFs into microcapsules and the need to consider GF release profile when designing capsule-based differentiation protocols. We note that timing of GF loading was part of the consideration for Hep^Imm/Imm^ microcapsules. We observed (see Fig. 3E) that >80% of pluripotency signals were released by day 5 of culture and chose to introduce endodermal signal, Nodal, at this time point. The immunofluorescence staining for Sox17 corroborated RT-PCR results, confirming higher level and frequency of expression of this endodermal marker for stem cells in Hep microcapsules compared to bare spheroids (Fig. 5E).

### 3.7 Modeling local GF concentrations experienced by encapsulated hPSCs

Improved maintenance and differentiation of encapsulated hPSCs may be explained in part by high local concentration of GFs in bioactive microcapsules. In order to assess local concentration of GF, we developed a COMSOL model that takes into account the affinity interactions with heparin and diffusion of GF molecules. The parameters used for constructing this model are provided in Table S2.

The results of modeling, presented in Fig. 6A, revealed that the local concentrations of FGF-2 and TGF-β1 were 1.6 and 1.4 times greater than the corresponding initial solution concentrations of these GFs and these higher levels stably persisted for at least 12 h after 1h loading phase. This prediction may be rationalized by high affinity of heparin for FGF-2 and TGF-β1 (1.2 nM and 59 nM, respectively [30]), which resulted in GF concentration in the hydrogel shell being 40 to 50-fold higher than in solution at the time of loading (see Fig. 6B). High affinity of the bioactive hydrogel shell also contributed to sustained levels of GFs inside the core of the microcapsule. Our modeling suggests that release behavior observed in Fig. 3 is governed/driven by exchanging of media and disturbing GF gradients, and that encapsulated stem cells are likely exposed to a constant concentration of GFs between exchanges of media.

**Fig. 6.**
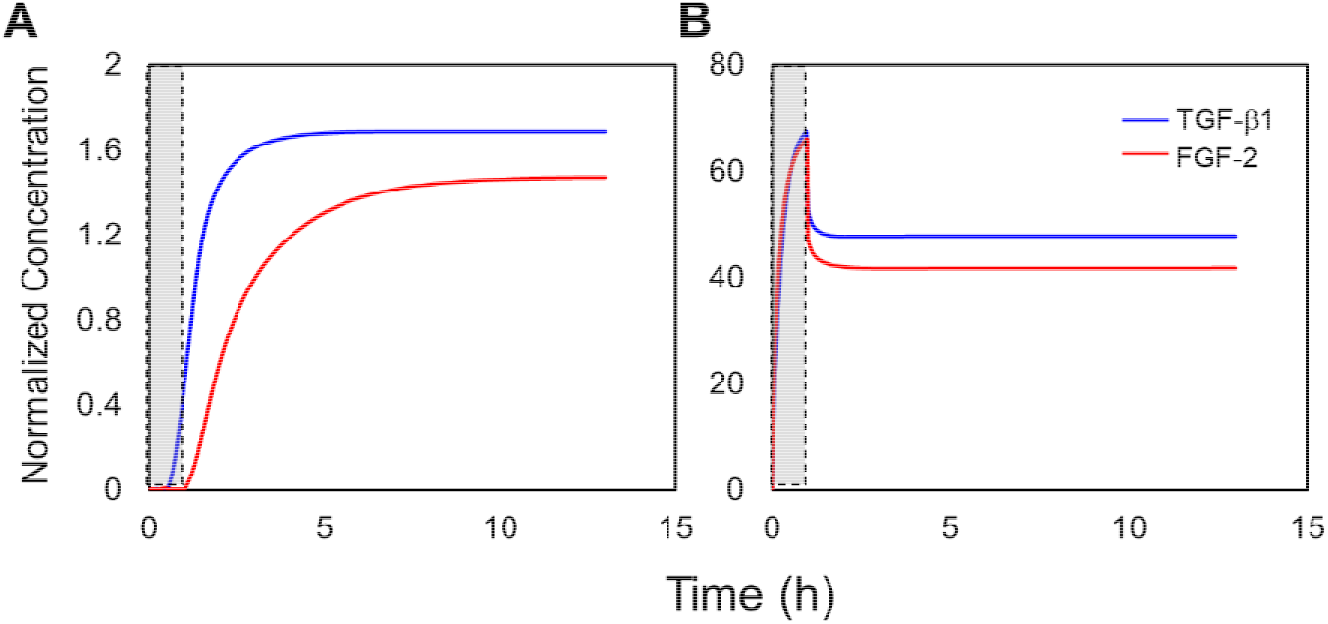
GF loading and release from a bioactive capsule. Average concentration of GF (red: FGF-2, blue: TGF-β1) was normalized to solution concentrations used for GF loading (Table S2) and plotted as a function of time (h) for (**A**) aqueous core and (**B**) hydrogel shell. The gray box indicates loading time of 1 h.

Taken together our modeling results suggest that stem cells are exposed to higher local concentrations of inductive signals compared to regular cultures. Furthermore, we expect the microcapsules to act as on-demand GF delivery depots where consumption of GFs by cells results in immediate and local release of new GFs. While we have not incorporated GF internalization by cells into the model, we expect the process of on-demand GF release from microcapsules to be far more dynamic than passive, diffusion-based delivery of GFs in standard culture systems. High local concentration of GFs inside the hydrogel shell (see Fig. 6B) may also contribute to juxtacrine signaling to the stem cells. Taken together, our modeling results may explain empirical observation of improved stem cell maintenance and differentiation in bioactive microcapsules.

## 4. Conclusion

This study investigated the use of bioactive heparin-containing microcapsules for maintenance and differentiation of hPSCs. The microcapsules contained an aqueous core, to facilitate stem cell aggregation and spheroid formation, as well as heparin hydrogel shell for loading and release of GFs. We characterized release profiles for key signals driving pluripotency maintenance and endodermal differentiation of hPSCs (FGF-2, TGF-β1, and Nodal), and demonstrated that all GFs exhibit sustained release from bioactive microcapsules over the course of 7 to 9 days. Importantly, we also demonstrated that one-time loading and local release of FGF-2 and TGF-β1 in Hep microcapsules induces a pluripotency state similar to or better than daily exposure to soluble GFs. We envision employing bioactive microcapsules for multi-step cultivation protocols where developmental cues are delivered to stem cells at specific temporal windows. To mimic such protocols, we demonstrated that the same Hep microcapsules may first be loaded with pluripotency signals and, after most of these signals have been released, may be reloaded with an endodermal cue, Nodal. Intriguingly, hPSCs differentiated inside bioactive Hep microcapsules expressed a higher level of endodermal markers compared to hPSCs exposed to soluble GFs. Overall, bioactive core-shell microcapsules represent an exciting new system for stem cell cultivation and offer the following benefits: 1) rapid and reproducible spheroid formation for 3D cultivation of hPSCs, 2) significant reduction in the amount of GFs needed for hPSC cultivation (>5 fold depending on the differentiation step), 3) potential improvements in stem cell phenotype/differentiation efficiency, and 4) scalable cultivation in stirred bioreactors as described by us recently [12]. In the future, we plan to employ bioactive core-shell microcapsules for pancreatic and hepatic differentiation of hPSCs.

## Acknowledgements

This study was supported in part by the grants from the Mayo Clinic Center for Regenerative Medicine, J.W. Kieckhefer Foundation, Al Nahyan Foundation, Regenerative Medicine Minnesota (RMM 101617 TR 004) and NIH (DK107255). Additional support was provided by an NIH Grant EB021911 to HB.

