## Supplementary Information for "Bioactive Hydrogel Microcapsules for Guiding Stem Cell Fate Decisions by Release and Reloading of Growth Factors"

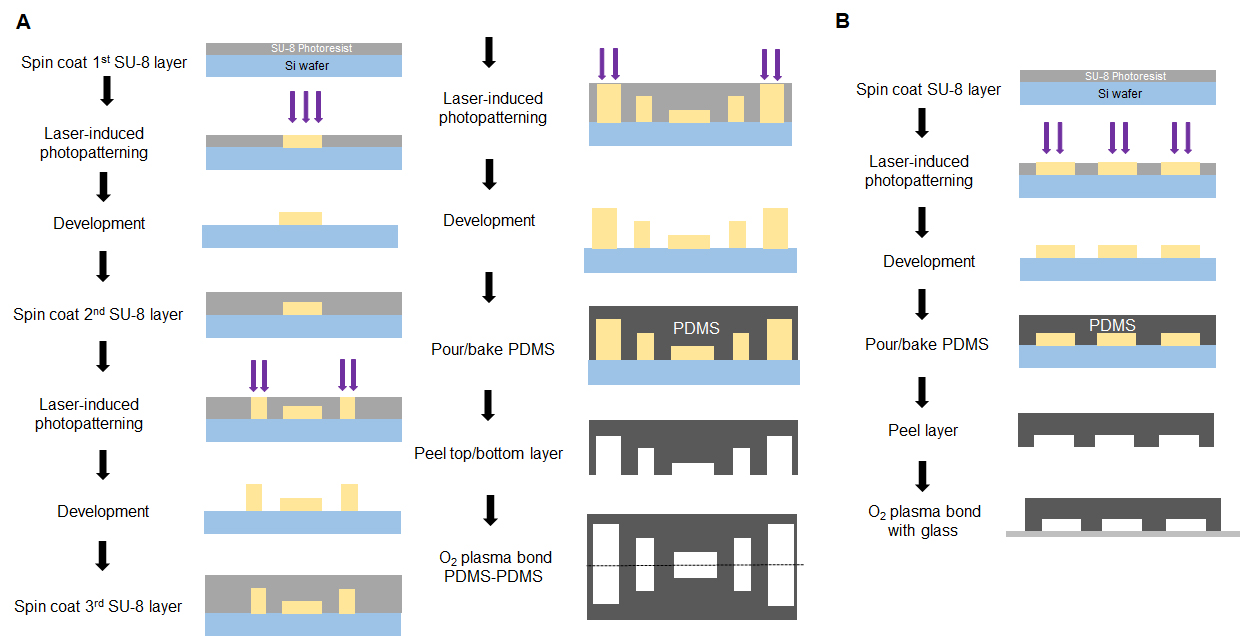

**Fig. S1.** Fabrication of microfluidic devices used for encapsulation. (A) Fabrication of the encapsulation device. Three layers of SU-8 photoresist were spin-coated, and photo patterned to generate core, shell, and oil channels with different heights. Top and bottom PDMS pieces were then aligned, and plasma bonded. (B) Fabrication of the filter device. One layer of SU-8 photoresist was spin-coated, and photo patterned to generate filter channels. PDMS piece was then plasma bonded to a glass slide.

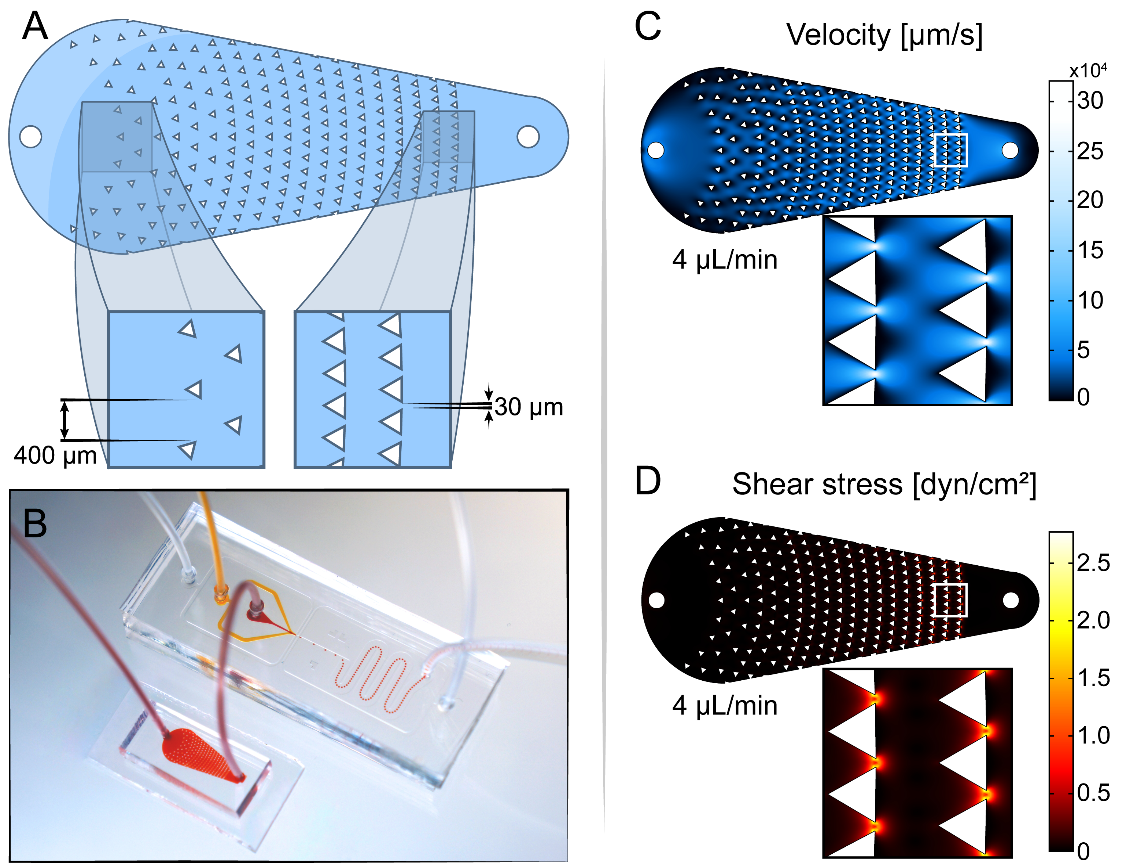

**Fig. S2.** (A) Design of the microfluidic filter device used to avoid large cell aggregates entering microencapsulation device. The device comprises an array of posts distributed in a sector form chamber. Posts separation is larger (400 µm) closer to the inlet, and decreases (30 µm) closer to the outlet. Larger cell aggregates are retained in the device, while cell aggregares with diameter <30 µm flow through the filter device and into the encapsulation device. (B) An image of the encapsulation system comprised of the filter and encapsulation devices connected in series. (C) Velocity (μm/s) profiles and (D) shear stress (dyn/cm^2^) profiles in the dissociation device at experimental flow rate, showing low shear stress levels even at the narrower areas, ensuring minimal cell harm while flowing through the filter device.

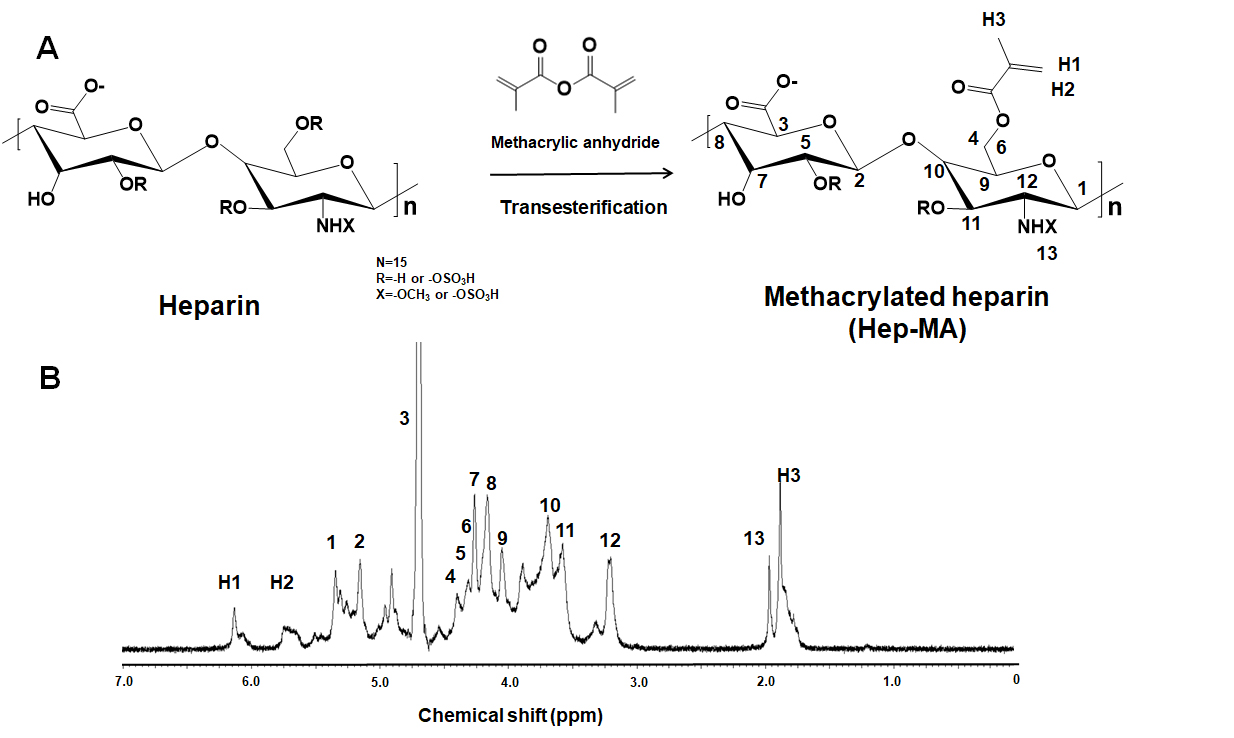

**Fig. S3.** (A) Synthesis of methacrylated heparin(Hep-MA) via transesterification reaction between heparin (HA) and methacrylic anhydride (MA). (B) The 1H-NMR spectra of Hep-MA. The degree of methacrylation was calculated by comparing the methacrylate peaks of MA at 5.6 and 6.1 ppm and the presenting protons 4–11 (3.4–4.6 ppm) and proton 12 (3.0–3.4 ppm) on the repeating disaccharide unit of heparin.

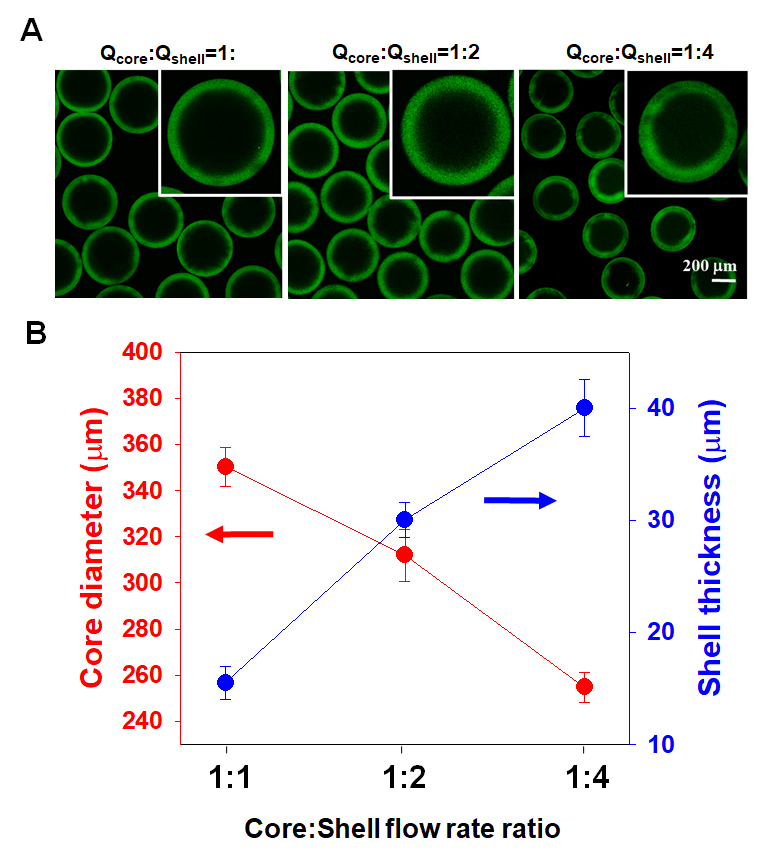

**Fig. S4.** Effects of the core and shell flow rates on the core size and shell thickness of the core-shell microcapsule. (A) Shell thickness (visualized with FITC-labeled PEG) was varied by modulating core/shell flow rate ratio. (B) The core diameter and shell thickness as a function of the core and shell flow rates ratio.

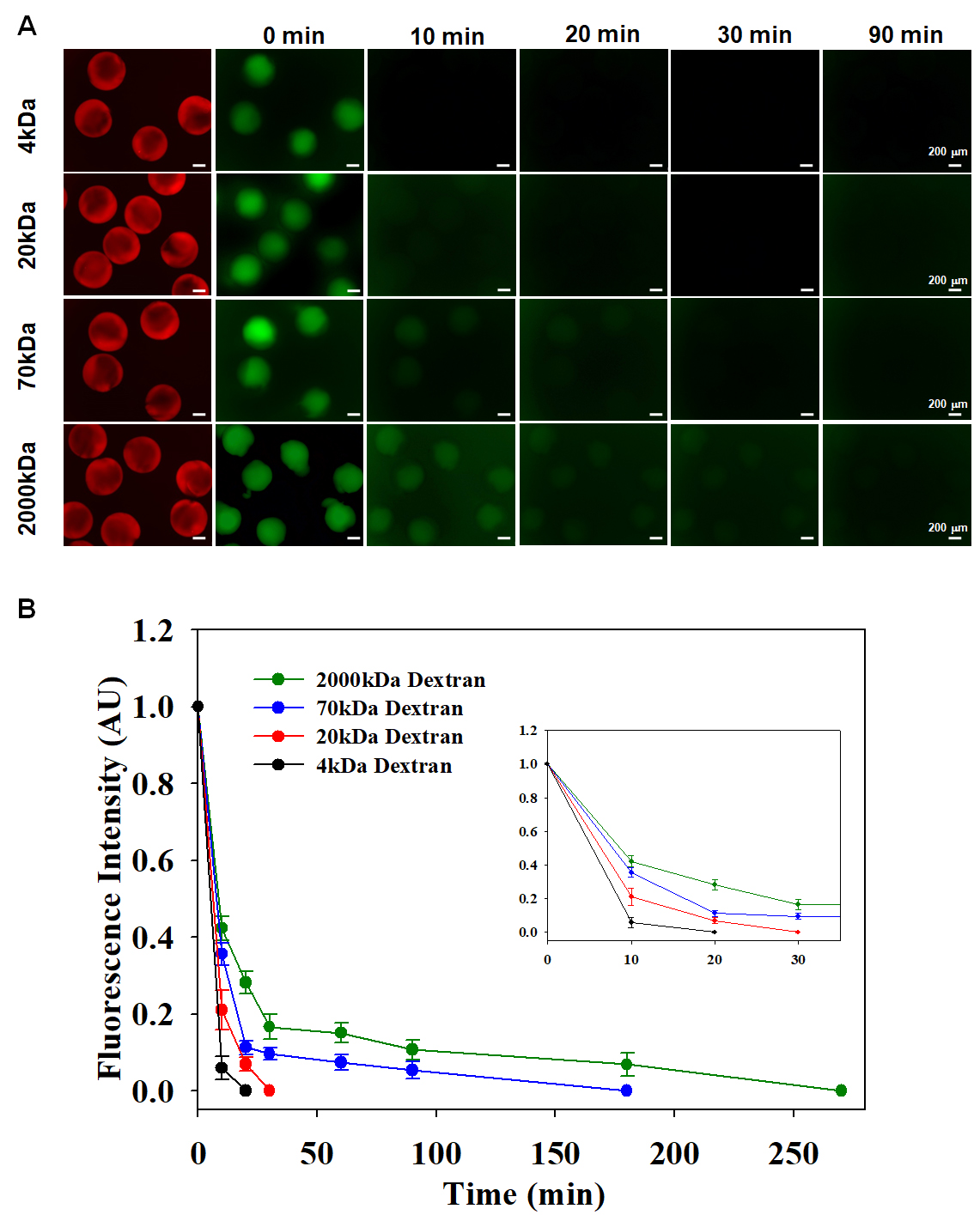

**Fig. S5.** Diffusion of model growth factors through the hydrogel shell of microcapsules. (A) Release of various FITC labeled dextran (MW 4, 20, 70, 2000 kDa) from heparin-based core-shell microcapsules. Rhodamin B-labeled PEG (red) is used to visualize the capsules. (B) Fluorescence intensity of capsules over time (n=7). The graph demonstrates that decrease in fluorescent intensity as a function of time and decreases diffusion rate as increase the molecular weight of dextran.

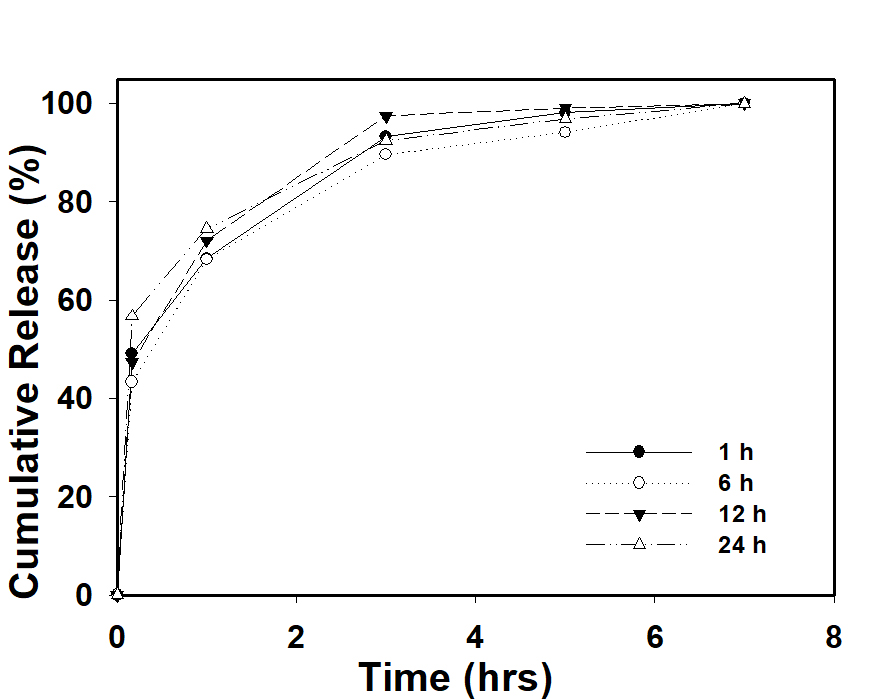

**Fig. S6.** FGF-2 release profile with different loading incubation times (1, 6, 12, and 24 h).

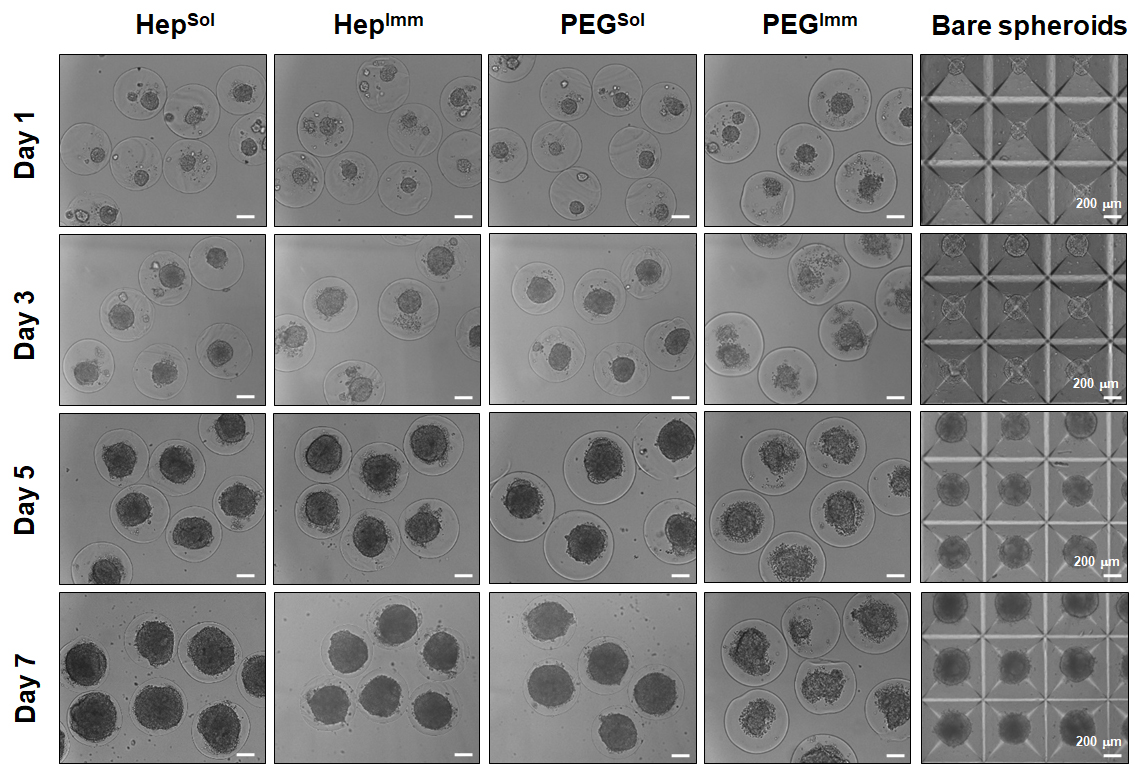

**Fig. S7.** Representative images of encapsulated HUES8 spheroids and bare spheroids cultured in different conditions over time.

**Table S1.** Sequences of primers used in qRT-PCR analysis.

| **Gene** | **Forward** | **Reverse** |
| --- | --- | --- |
| **GAPDH** | **TGTTGCCATCAATGACCCCTT** | **CTCCACGACGTACTCAGCG** |
| **SOX2** | **TCAGGAGTTGTCAAGGCAGAGAAG** | **GCCGCCGCCGATATTGTTATTAT** |
| **OCT4** | **GATCACCCTGGGATATACAC** | **GCTTTGCATATCTCCTGAAG** |
| **NANOG** | **CCGGTCAAGAAACAGAAGACCAGA** | **CCATTGCTATTCTTCGGCCAGTTG** |
| **GATA4** | **CATCAAGACGGAGCCTGGCC** | **TGACTGTCGGCCAAGACCAG** |
| **CXCR4** | **CTTCATCTTTGCCAACGTCAG** | **GGACAGGATGACAATACCAGG** |
| **SOX17** | **GGCGCAGCAGAATCCAGA** | **CCACGACTTGCCCAGCAT** |
| **FOXA2** | **CGAGTTAAAGTATGCTGGG** | **CATGTACGTGTTCATGCC** |

**Table S2.** lists all the parameters used in the simulations.

| Parameter | Value |
| --- | --- |
| Overall diameter of the microcapsule | 400 µm |
| Shell thickness | 15 µm |
| Cell spheroid diameter | 300 µm |
| Diameter of fluid layer surrounding capsule | 1.04 mm |
| Loading concentration of FGF-2 | 6.06 nM |
| Loading concentration of TGF-β1 | 0.156 nM |
| Loading duration | 1 h |
| Total heparin concentration in shell | 3.15 mM |
| Diffusivity of FGF-2 in medium | 5.6 × 10^-11^ [m^2^/sec]  (in water, [1]) |
| Diffusivity of FGF-2 in shell | 4.425 × 10^-12^ [m^2^/sec]  (estimated in this work) |
| Diffusivity of TGF-β1 in medium | 3.3 × 10^-11^ [m^2^/sec]  (in water, [1]) |
| Diffusivity of TGF-β1 in shell | 2.27 × 10^-12^ [m^2^/sec]  (estimated in this work) |
| $k_{FGF-2}^{ON}$ | 2.4 × 10^6^ [1/Ms] [1] |
| $k_{FGF-2}^{OFF}$ | 2.0 × 10^-3^ [1/s] [1] |
| $k_{TGF-\beta1}^{ON}$ | 1.0 × 10^5^ [1/Ms] [1] |
| $k_{TGF-\beta1}^{OFF}$ | 9.2 × 10^-3^ [1/s] [1] |

*Characterization of Shell Diffusive Properties*

1-D unsteady radial transport equation for the dextran in the core is given by $\frac{\partial C}{\partial t}=\frac{D}{r^{2}}\frac{\partial}{\partial r}\left( r^{2}\frac{\partial C}{\partial r} \right)$. For the release studies, the initial and boundary conditions are:$at t=0, C=C_{0}, at r=0, \frac{\partial C}{\partial r}=0, at r=R,-D\frac{\partial C}{\partial r}=kC$. Here, R is the radius of the core, k is the permeability of the dextran molecule in the shell. The above equations can be non-dimensionalized to obtain $\frac{\partial C'}{\partial t'}=\frac{1}{{r'}^{2}}\frac{\partial}{\partial r'}\left( {r'}^{2}\frac{\partial C'}{\partial r'} \right)$ with the initial and boundary conditions: $at t'=0, C'=1, at r'=0, \frac{\partial C^{'}}{\partial r^{'}}=0, at r'=1,-\frac{\partial C^{'}}{\partial r^{'}}=PeC'$. Here, $C^{'}=\frac{C}{C_{0}}, r^{'}=\frac{r}{R}, t^{'}=\frac{Dt}{R^{2}}, and Pe=\frac{kR}{D}$. The dimensionless parameter, Pe, was adjusted to fit the model data (C/C_0_) to experimental data (I/I_0_) as described in Methods. The estimated permeability values for various dextran molecules are shown in the following table. Diffusivity values can be obtained by assuming that the solubilities of dextran molecule in the core aqueous solution and in the shell are similar.

**Table S3**. Diffusivity in shell regarding molecular weight of dextran.

| **Mw of dextran**  **[kDa]** | **Diffusivity of dextran [m^2^/s]** | **Permeability**  **[m/s]** | **Diffusivity in Shell**  **[m^2^/s]** |
| --- | --- | --- | --- |
| 4 | 102 × 10^-12^ | 3.74 × 10^-7^ | 5.9 × 10^-12^ |
| 20 | 44 × 10^-12^ | 2.2 × 10^-7^ | 3.44 × 10^-12^ |
| 70 | 33 × 10^-12^ | 1.44 × 10^-7^ | 2.27 × 10^-12^ |
| 2000 | 9 × 10^-12^ | 1.96 × 10^-7^ | 3.08 × 10^-12^ |
